# Design and Validation of a Hybrid Stimulus fMRI Paradigm for Simultaneous Retinotopy and Mapping of Reading-Selective Visual Cortex

**DOI:** 10.64898/2026.07.20.739334

**Authors:** Miguel Martinez-Zaldivar, Yongning Lei, Garikoitz Lerma-Usabiaga

**Affiliations:** BCBL. Basque Center on Cognition, Brain and Language, Donostia-San Sebastián, Basque Country, Spain; IKERBASQUE - Basque Foundation for Science, Basque Country, Spain

**Keywords:** Reading, vision, MRI, vOTRC, VWFA, pRF, Interactive Account, Word-Center paradigm

## Abstract

Reading selectively recruits specialized cortical patches within the left ventral occipitotemporal cortex (vOTC), such as the visual word form area (VWFA). Under the framework of the interactive account of word recognition, these neural responses emerge from dynamic, bidirectional loops where top-down linguistic predictions continuously modulate and constrain bottom-up sensory visual inputs. Consequently, identifying and isolating these top-down cognitive signals fundamentally requires a precise characterization of the underlying low-level sensory baseline.

This dense-sampling fMRI study (11 right-handed adults, 10 sessions each) systematically addresses the spatial and functional architecture of the reading network using population receptive field (pRF) modeling and functional localizers (fLoc). First, we conduct a rigorous group- and subject-level replication of stimulus-dependent pRF eccentricity shifts (words and false fonts versus checkers) observed in previous research. To overcome known barriers to individual parameter stability, we systematically manipulate and evaluate stimulus Frequency, bar Width, and element Size (FWS) across eight sessions per subject to isolate the precise factors driving test-retest reliability. Second, we design and validate a novel Word-Center (WC) paradigm acquired across one/two sessions per participant. This hybrid paradigm was designed to decouple moving spatial mapping carriers (checkerboard bars) from central reading processes (word or false-font streams in the central fixation location). Functional data were denoised with NORDIC, preprocessed via fMRIPrep, and projected to the cortical surface for analysis via Nilearn GLMs to model word-responsive regions and mrVista to obtain pRF estimates.

Individual-level spatial maps replicated previous results in approximately 70% of subjects. The lack of replication in the remaining subjects, whether driven by methodological factors or inherent individual differences, highlights ongoing baseline mapping challenges. Optimizing this mapping is sensitive to stimulus carrier type and specific FWS configurations. Furthermore, the hybrid WC paradigm successfully demonstrates stimulus equivalence, concurrently yielding robust reading-selective functional localization and reliable retinotopic estimates. By breaking the spatial-lexical confound of traditional mapping protocols, this paradigm provides an innovative framework to separate bottom-up sensory sweeps from top-down central linguistic processing. This vision-centric framework serves as a methodological baseline toward a comprehensive, multi-signal understanding of the human reading hierarchy.

## Introduction

Reading is a complex cognitive milestone that demands the orchestration of sensory, perceptual, and linguistic systems within a fraction of a second (Caffarra et al., 2021; Yeatman & White, 2021). Although fluent reading feels effortless, it represents a notable computational challenge for the brain: arbitrary visual symbols (graphemes) must be mapped onto corresponding phonological and semantic representations with speed and precision. A long-standing question in cognitive neuroscience is how the visual system accommodates these unique, language-specific demands. While mature reading relies on specialized cortical regions within the ventral stream (Cohen et al., 2000), understanding how these regions process linguistic text fundamentally requires a thorough characterization of the low-level visual inputs that feed into them. Consequently, although the empirical methods, results, and analyses of this study are extensively rooted in vision and retinotopic mapping, this vision-centric approach is a deliberate and mandatory stepping stone. To ultimately decode the neural mechanisms of reading, we must first establish a highly precise, low-level sensory baseline. By fully mapping how the visual cortex represents space and handles basic stimulus characteristics we plan to isolate and understand the top-down cognitive signals that transform a visual shape into a meaningful word.

### The Vision-Reading Nexus: The Interactive Account

To understand the relationship between visual perception and language, this study adopts the framework of the interactive account of word recognition (Price & Devlin, 2011). Historically, models of reading often posited a strict, feedforward hierarchy, where basic visual features are extracted in primary visual cortex (V1) and sequentially integrated into complex shapes until reaching a dedicated visual word module in the vOTC (Dehaene & Cohen, 2011).

However, the Interactive Account challenges this passive view, conceptualizing the reading network as a dynamic, bidirectional system. In this model, neural responses within reading-selective regions, collectively termed the Ventral Occipito-Temporal Reading Circuitry (vOTRC), do not represent a simple feedforward accumulation of visual inputs. Instead, they emerge from the continuous interaction between bottom-up sensory signals originating from occipital visual cortex and top-down cognitive or linguistic predictions originating from perisylvian language areas (Price & Devlin, 2011; Saygin et al., 2016). Under this interactive framework, top-down predictions (e.g., phonological and semantic expectations) actively modulate and constrain the incoming visual data. In order to identify, measure, and isolate these top-down linguistic signals, we should first possess an understanding of the bottom-up sensory baseline. Without a mapping of the underlying sensory processing, variations caused by top-down cognitive modulations could be misattributed to basic visual differences, or vice-versa. Therefore, a deep dive into the retinotopic and spatial properties of the visual cortex is a logical requirement to unravel the interactive nature of reading.

### Functional Segregation and Sensory-Cognitive Integration in the vOTRC

The necessity of a precise sensory baseline becomes even more apparent when examining the functional landscape of the left ventral temporal cortex. Rather than acting as a single, homogeneous block, neuroimaging studies reveal that the vOTRC is segregated into distinct, specialized subregions organized along a posterior-to-anterior gradient of abstraction (Lerma-Usabiaga et al., 2018; Yablonski et al., 2024). At successive stages of the ventral stream, sensitivity to abstract lexical features increases while sensitivity to low-level visual properties decreases (Li et al., 2024; Dehaene et al., 2005). Specifically, the vOTRC is functionally divided into at least two distinct anatomical sites (Figure 1): the posterior occipitotemporal sulcus (pOTS) is primarily engaged in parallel visual feature extraction and letter-shape processing (White et al., 2019; Caffarra et al., 2021), receiving dense structural inputs from attention-related parietal networks via the Vertical Occipital Fasciculus (VOF; Yablonski et al., 2024; Weiner et al., 2017b). Moving anteriorly, the mid-occipitotemporal sulcus (mOTS) serves as a processing bottleneck, interfacing directly with frontal language networks via the Arcuate Fasciculus (AF) to mediate lexical access (Lerma-Usabiaga et al., 2018; White et al., 2019; Yablonski et al., 2024). These functional differences are further supported by the underlying microarchitecture; the pOTS resides within the cytoarchitectonic area FG2, while the mOTS is localized to FG4 (Lerma-Usabiaga et al., 2018; Weiner et al., 2017a; Caffarra et al., 2021).

**Figure 1.**
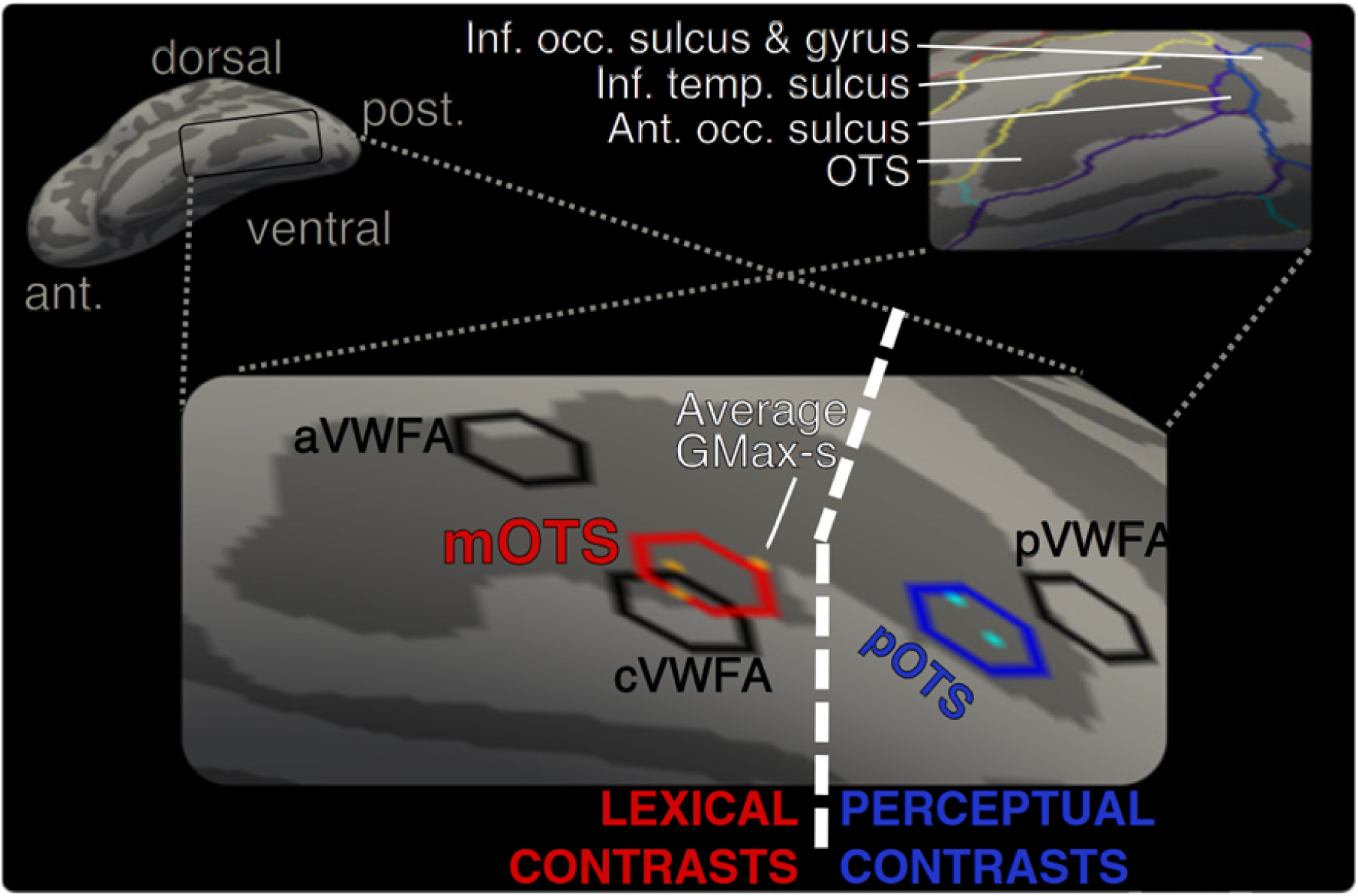
Functional segregation of the vOTRC. Reprinted from “Converging evidence for functional and structural segregation within the left ventral occipitotemporal cortex in reading”, by G. Lerma-Usabiaga, M. Carreiras, & P. M. Paz-Alonso, 2018, Proceedings of the National Academy of Sciences of the United States of America, 115(42), E9981–E9990, Figure 1, p. E9982 (https://doi.org/10.1073/pnas.1803003115). © 2018 National Academy of Sciences. Published under the PNAS License; reprinted with permission.

Because this transition from low-level visual shapes to abstract linguistic entities is distributed across closely adjacent, heterogeneous patches, mapping the fine-grained visual properties of this hierarchy is crucial. To understand how the brain transitions from seeing a shape to reading a word, we must precisely map how these subregions respond to spatial coordinates, establishing exactly where sensory dominance ends and cognitive modulation begins. Other research has proposed alternative segmentations of the reading regions, ranging from the classic single-module Visual Word Form Area (VWFA) to more extensive multi-patch systems. A further and more precise segmentation of these reading regions, particularly one that moves toward individual-specific functional boundaries, would be highly beneficial to resolve these diverse accounts and provide a more accurate map of the sensory-cognitive transition

### Retinotopic organization and the Population Receptive Fields

A fundamental organizing principle of the human visual system is retinotopy, the process by which the spatial layout of the retina is preserved as an orderly map of visual space on the cortical surface (Wandell & Winawer, 2011). To precisely quantify these visual responses, contemporary research utilizes population receptive field (pRF) modeling (Dumoulin & Wandell, 2008). In this technique, spatial tuning properties of neuronal populations are recovered by measuring how the fMRI response varies as a stimulus aperture moves across the visual field over time, sequentially activating different regions (Figure 2, panelA).

**Figure 2.**
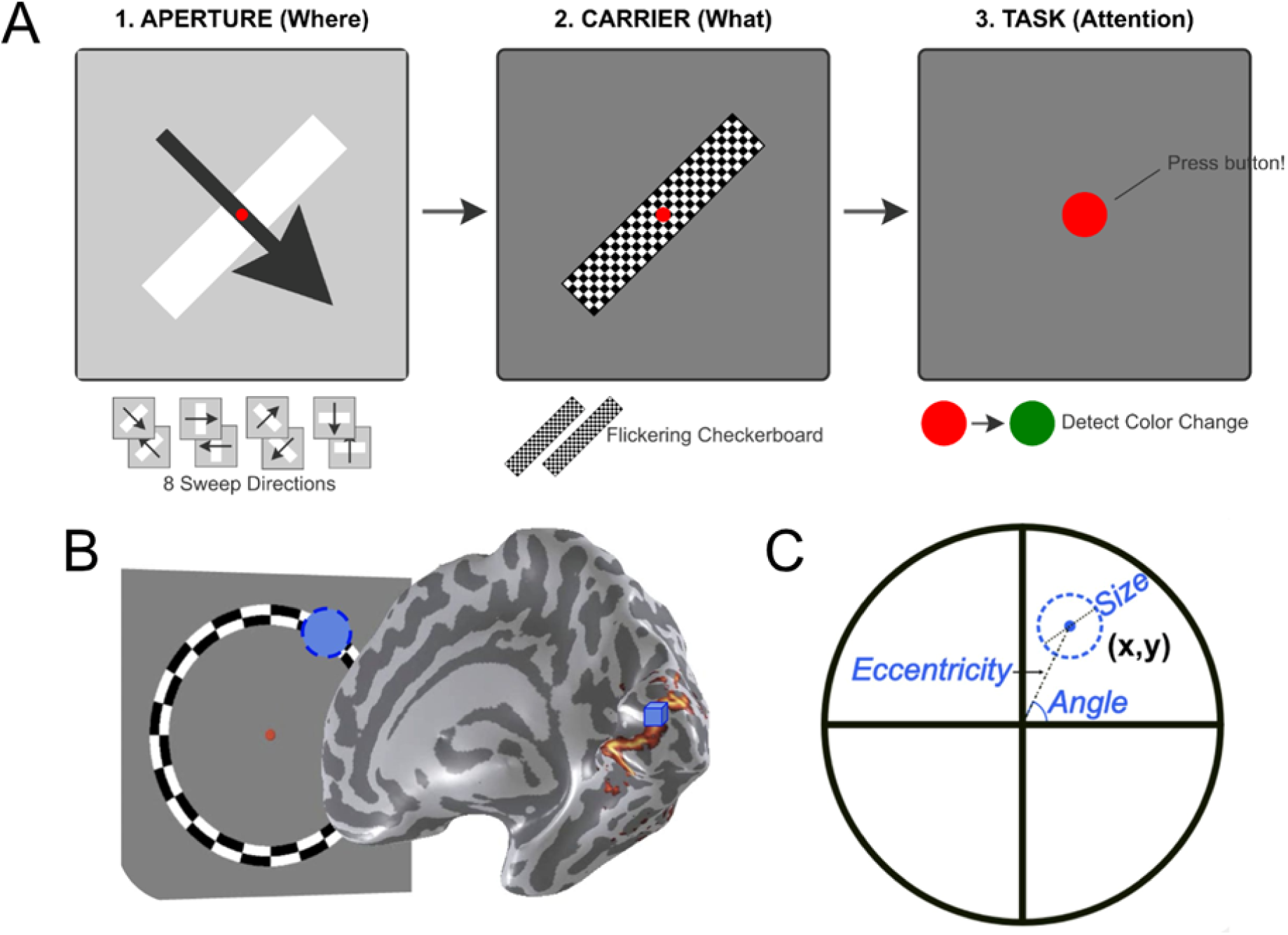
pRF modeling. **A.** Visual stimuli typically presented to an individual in a retinotopic mapping experiment. 1. shows an aperture that defines where the stimulus is at any moment in time. This aperture traverses the visual field, revealing the stimulus/carrier from the upper left to the lower right along the diagonal. At the bottom, all directions the aperture traverses are shown to cover the entire visual field and minimize biases due to the hemodynamic response delay. 2. An example of a carrier, the stimulus that is revealed by the aperture, used in retinotopic mapping experiments. 3. To minimize eye movements and ensure participants’ wakefulness, retinotopic mapping visual stimuli are paired with a color-change detection task, during which participants press a button whenever the fixation dot changes color. **B.** Retinotopic response in the brain. The objective is to find which part of the visual field is computed by each voxel/vertex in the visual cortex. **C.** pRF parameter estimate visualization. The circle represents the visual field of a participant, and the center (x,y or eccentricity, angle) and size represent the pRF.

A pRF is a computational model that summarizes the fMRI signal of a single voxel by estimating the specific region of the visual field to which its underlying neuronal population is sensitive (Figure 2, panel B): it provides a quantitative description of every voxel’s center coordinates (x,y) and its receptive field size (σ), allowing retinotopic organization mapping and measurement of cortical visual-field coverage (Dumoulin & Wandell, 2008). A key measure used in pRF modeling is eccentricity, which refers to the distance in degrees between the point of fixation (fovea) and a stimulus location in the visual field (Figure 2, panel C). Eccentricity values are used to determine whether a cortical site responds preferentially to foveal or peripheral visual inputs (Dumoulin & Wandell, 2008; Wandell & Winawer, 2011). Beyond mapping retinotopic organization, pRF modeling has established that receptive field sizes systematically grow as a function of both stimulus eccentricity and hierarchical distance from V1 (Yeatman & White, 2021). Consequently, pRFs are smallest in V1 and become progressively larger as signals advance through extrastriate areas toward high-level visual regions (Smith et al., 2001).

A defining characteristic of high-level visual cortex revealed by pRF modeling is eccentricity bias, an organizing principle where cortical regions are anatomically segregated based on their sensitivity to central versus peripheral visual field locations (Wandell & Winawer, 2011). Under this framework, regions specialized for categories requiring high-resolution vision, such as faces and written words, are associated with center-biased (foveal) representations (Hasson et al., 2002; Nordt et al., 2019). Conversely, regions involved in large-scale feature integration, such as place-selective areas in the collateral sulcus, show a bias toward the visual periphery (Hasson et al., 2002).

The vOTRC exhibits a high foveal specialization, with pRF centers concentrated within the central visual field (Dehaene & Cohen, 2011; Yeatman & White, 2021). This high-resolution focus is functionally necessary for resolving the small and complex configurations of line junctions that define written words (Dehaene & Cohen, 2011). While pRF modeling has successfully mapped the spatial sensitivity, recent estimates have revealed a new complication: the resulting spatial maps appear to be stimulus-dependent, shifting their positional estimates when measured with words compared to spatially matched checkerboard carriers (Lerma-Usabiaga et al., 2021; Le et al., 2017). This dependency presents a major methodological challenge for purely sensory models of vision, as it suggests that signals in the reading circuitry are not fixed reflections of space, but instead represent a complex integration of bottom-up sensory information and top-down cognitive signals.

### Stimulus-Dependency and the Methodological Challenge of Replicability

The intersection of vision and reading is brought into sharp focus by a critical discovery: spatial maps in high-level visual regions are not static, but are influenced by the type of stimulus used to measure them (Le et al., 2017; Lerma-Usabiaga et al., 2021). In a recent study, Lerma-Usabiaga et al. (2021) demonstrated that word-like stimuli elicit pRF center estimates that shift closer to the fovea (by up to 3–4 degrees) compared to standard checkerboard carriers. This effect became progressively more pronounced in successively higher-order ventral ROIs. It was observed in approximately 80% of the participants, whereas 10% showed no measurable effect and the remaining 10% exhibited the opposite pattern.

This stimulus-dependency reveals a functional dissociation between early visual maps and the reading network. In the VO-1 ventral map, responses to words and false fonts are equivalent, and both differ from checkers, indicating a purely sensory grouping based on visual line-junction characteristics. However, upon entering the vOTRC, the pattern reverses: false fonts and checkers evoke similar responses, while words evoke a highly distinct, foveal pattern (Figure 3). This finding indicates that the vOTRC integrates a top-down cognitive language signal that isolates true lexical items from visually identical non-words (Lerma-Usabiaga et al., 2021).

**Figure 3.**
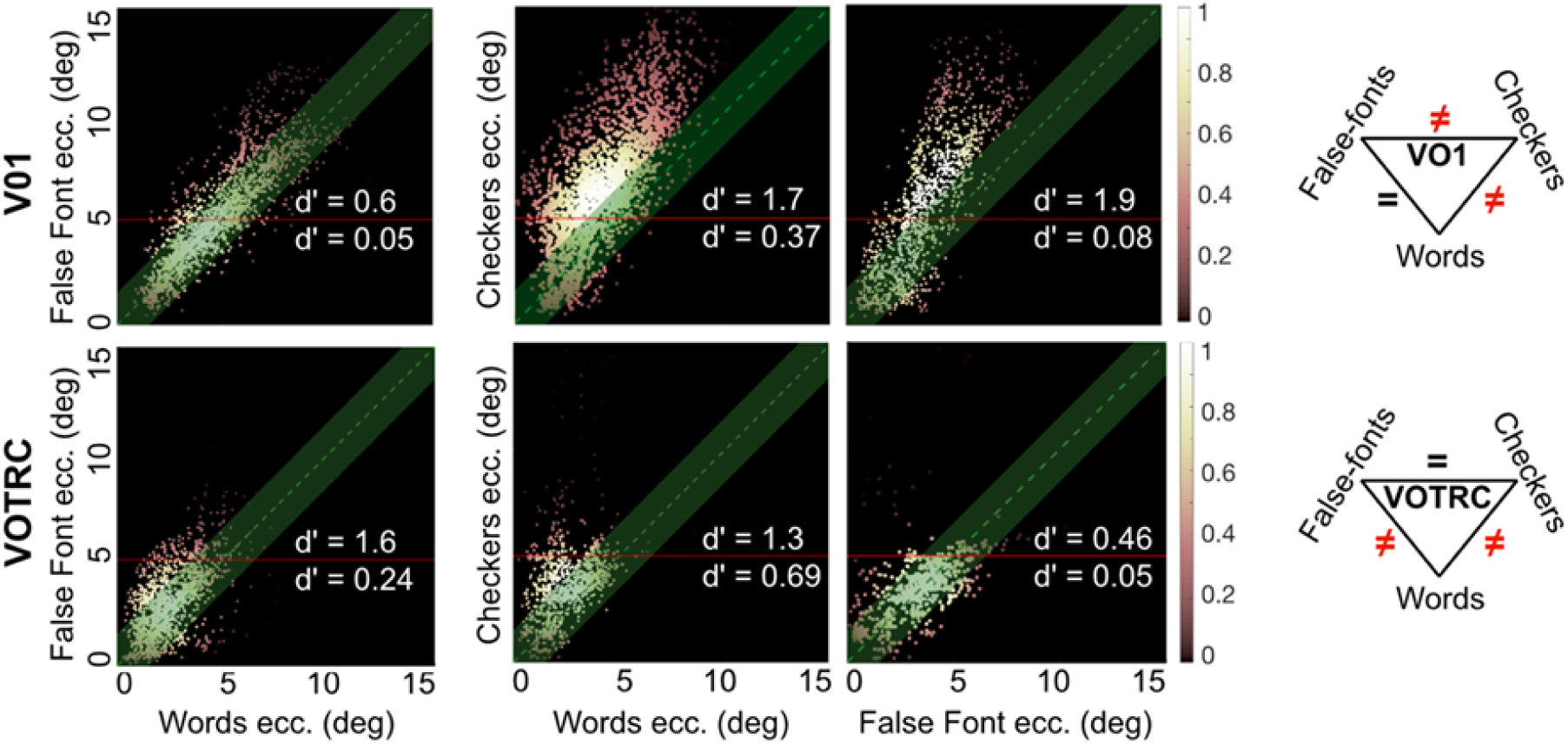
Comparison of the pRF centers for English words, False-Fonts, and Checkers. Reprinted from “Interpreting sensory and cognitive signals in the cortical reading network”, by G. Lerma-Usabiaga, R. Le, C. Gafni, M. Ben-Shachar, B. A. Wandell, 2021, BioRxiv, Figure 3 (https://doi.org/10.1101/2021.09.14.460238). Reprinted with permission.

However, translating these group-level findings into a robust, subject-specific model presents an intense methodological challenge. Because word-selective cortical patches are tiny and highly variable across individuals, group-level averages frequently mask the true functional boundaries of the circuitry (Glezer & Riesenhuber, 2013). Capturing these stimulus-dependent spatial shifts at the individual subject level is difficult, and initial replication efforts often face significant obstacles. This fragility highlights a major gap in our understanding: we do not yet know how specific parameters of the experimental stimulus, such as flickering frequency, bar width, and element size (FWS), affect the reliability and replicability of pRF maps. Systematically testing how these FWS factors influence individual subject maps is a crucial prerequisite to building a stable sensory baseline.

### Current study

The present study is explicitly designed to address these fundamental challenges, bridging the gap between low-level visual space and high-level reading architectures through two primary objectives. First, we conduct a rigorous individual subject-level replication of the stimulus-dependent pRF effects described by Lerma-Usabiaga et al. (2021). We systematically manipulate and analyze the specific configurations of the stimulus, examining how variations in frequency, width, and size (FWS) impact test-retest reliability and individual parameter stability. This comprehensive evaluation provides the empirical groundwork to identify why initial replication attempts fail and how to optimize parameters to successfully capture these effects in single subjects.

Second, we design and validate a new experimental paradigm: the Word-Center (WC) paradigm. Traditional retinotopic protocols interweave the carrier (e.g., words or checkers) directly inside a moving bar, meaning that spatial position and lexicality are confounded. To disentangle these influences, the WC paradigm presents a standard, non-lexical moving checkerboard bar across the visual field while simultaneously presenting a continuous stream of text or false fonts at the central fixation point. While the WC paradigm is not fully orthogonal (as central attention and peripheral sweep still interact) it represents a major conceptual and methodological advance. By decoupling the moving spatial probe from the primary reading stimulus, this hybrid paradigm provides an innovative framework to cleanly separate bottom-up sensory sweeps from top-down central cognitive processing in future studies. Ultimately, by establishing a sensory baseline and validating this novel paradigm within subject-specific reading regions, this study provides a framework toward our ultimate goal: a comprehensive, multi-signal understanding of the human reading network.

## Methods

### Participants

Ten young adults (6 females; mean age = 29.2 ± 4.5 years) and an additional 11th control subject (male; 48 years old) participated in the study. Each participant completed 10 sessions with a minimum intersession interval of 3 days and a maximum of 2 weeks. All participants were right-handed healthy adults, with no history of psychiatric, neurological, attention, or learning disorders, and with normal or corrected-to-normal vision, and all of them gave written informed consent. The experiment was approved by the BCBL Ethics Review Board and complied with the guidelines of the Helsinki Declaration.

### Materials

Participants completed ten runs of a block-design fMRI experiment (5.5 minutes per run) adapted from Lerma-Usabiaga et al. (2018) and the fLoc functional localizer (Stigliani et al., 2015), aimed at identifying category-selective responses in the visual cortex. Each run included 51 randomized 6-s blocks, consisting of 42 active and 9 baseline blocks. The activation blocks contained 6 stimulus categories: words (RW; Figure 4, panel A1), false-fonts (FF; Figure 4, panel A2), consonant strings (CS; Figure 4, panel A3), scrambled words (SC; Figure 4A4), body/limb images (Figure 4, panel A5), and adult/infant face images (Figure 4, panel A6). Two counterbalanced stimulus sets containing 80 examples per category were assigned across participants. The experiment was counterbalanced across participants using these two sets. Stimuli were presented for 400 ms with a 100 ms inter-stimulus interval. Participants performed a one-back task throughout the experiment where they were asked to attend to the stimuli and respond once they detected that two sequential images were identical. Before entering the scanner, participants practiced a behavioural version of the experiment to familiarize themselves with the task and stimulus set.

**Figure 4.**
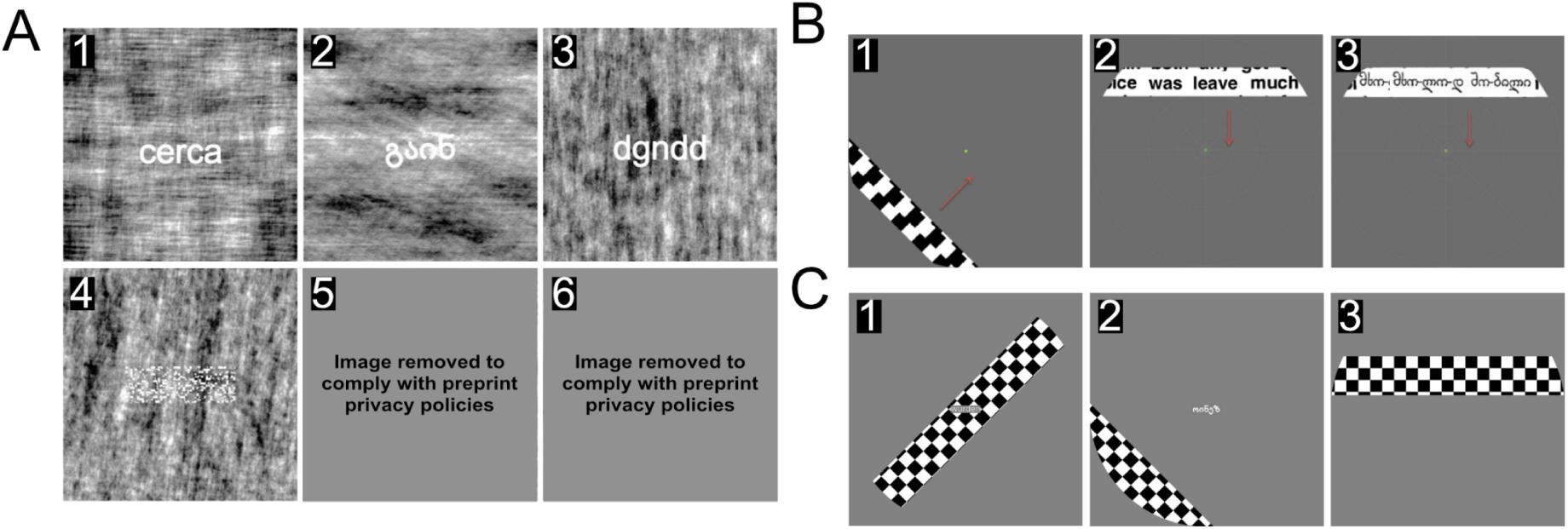
Examples of the three stimuli paradigms. **A.** fLOC stimulus example. A1. Words. A2. False-fonts. A3. Consonant strings. A4. Scrambled words. A5. Body/limb images (covered due to privacy policies). A6. Adult/infant face images (covered due to privacy policies). **B.** Retinotopy stimulus example: B1. Checkerboards. B2. Words. B3. False-fonts. **C.** Word-center stimulus example: C1. fixRW. C2. fixFF. C3. fixRWblock (blank interval), where fix stands for fixation as the words and false fonts were shown in the central fixation location continuously, except in the block case where they were shown in alternating on-off blocks.

Additionally, participants completed 6 retinotopy runs (5 minutes per run). Retinotopic-mapping stimuli were presented within moving bars (8 directions: left-to-right, right-to-left, up-down, down-up, and the four diagonals) while participants were asked to fix their gaze on the center of the screen. There were three different types of stimuli within the bars. Classic checkerboard stimuli (CB; Figure 4, panel B1), a variant with word stimuli (RW; Figure 4, panel B2), and a variant with false-font stimuli (FF; Figure 4, panel B3). For each session, participants were shown two runs of each stimulus condition. Across sessions we tested 3 flickering frequencies of alternating checkerboards or word movements (2, 4 and 8 Hz), 2 bar widths of (3° and 4° of visual angle), and 4 element sizes (25, 26, 50 and 100 pixels, corresponding to 0.44°, 0.46°, 0.88° and 1.76° of visual angle). The element size corresponds to the letter height for the RW and FF and to the side of a chequer square in CB. Eventually, we decided not to manipulate the bar widths, so the 4° sessions were excluded from analysis in this experiment (see Limitations). This results in a total of 9 different combinations: 3 flickering frequencies (2, 4, 8) x 1 bar width (3) x 3 element sizes (25, 50, 100). We summarize the manipulation throughout this work as FWS (Frequency, Width, Size). See Figure 5 for a subject x session table showing the FWS structure of the gathered data.

**Figure 5.**
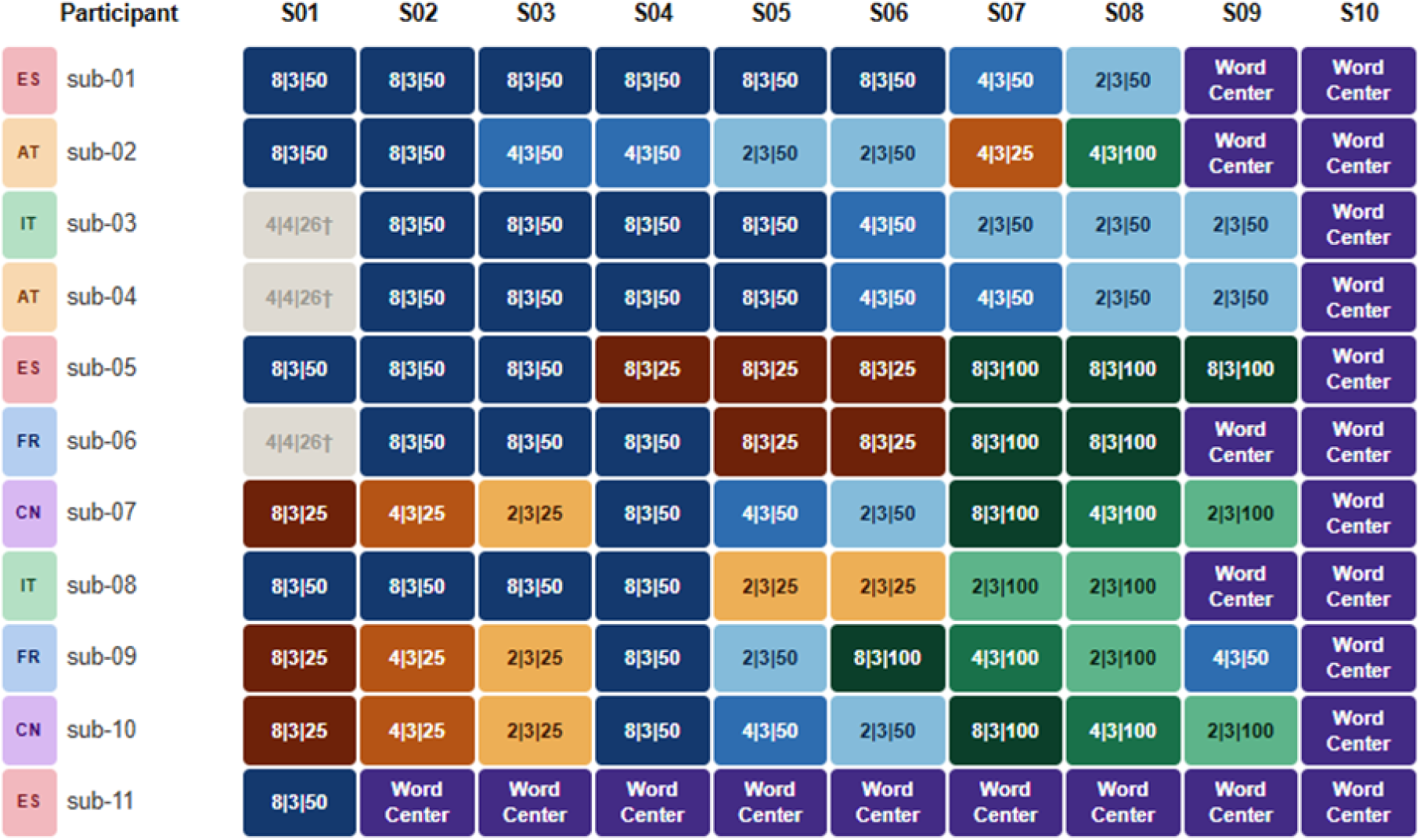
Session configuration across participants. Each session has three different values, illustrating the FWS (Frequency|Width|Size) configurations. The Word-Center sessions used the 8|3|50 configuration with the WC stimuli. The words in each participant were shown in a different language. *ES: Spanish; AT: German; IT: Italian; FR: French; CN: Mandarin Chinese*.

In the last session (last two sessions of 4 participants and last nine sessions of the subject 11), instead of the retinotopic runs with classical stimuli, participants completed the new experimental Word-Center paradigm (Figure 4, panel C). This new paradigm was developed as a sensory-cognitive hybrid to simultaneously engage bottom-up sensory and top-down linguistic processing. This approach allows us to evaluate whether the paradigm can effectively capture visual signals comparable to classical retinotopy while concurrently isolating the cognitive signals comparable to fLoc that characterize the functional specialization of the vOTRC. In all 6 runs of this paradigm, moving bars with a checkerboard carrier travelled across the screen while stimuli appeared and disappeared in the middle of the screen at a constant frequency. Three stimulus conditions were designed, differing in the type and temporal structure of the central stimuli. A continuous stream of real words (fixRW; Figure 4, panel C1), a continuous stream of false-fonts (fixFF; Figure 4, panel C2), and real words presented in alternating 10-second blocks of presentation and blank intervals (fixRWblock; Figure 4, panel C3).

### Data acquisition and preprocessing

Participants were scanned in a 3T Siemens Prisma Fit whole-body MRI scanner (Siemens Medical Solutions), using a 64-channel head coil. Headphones (MR Confon) were used to dampen background scanner noise and to enable communication with experimenters while in the scanner. To limit head movement, the area between participants’ heads and the coil was padded with foam, and participants were asked to remain as still as possible.

#### Structural imaging

T1-weighted volume was acquired with the MP2RAGE method with 1 mm3 isotropic voxel size and the following protocol parameters: TR/TE = 5000/2.98 ms, TI1/FA1 = 700 ms/4 deg, TI2/FA2 = 2500 ms/5 deg, bandwidth = 240 Hz/Pix, echo spacing = 7.14 ms, FOV = 256 × 240 mm with 176 slices, with R = 3 acceleration in the primary phase encoding direction (32 reference lines) and online GRAPPA image reconstruction, for a total acquisition time of 8 min 22 s per volume (Marques et al., 2010). A T2-weighted scan was also acquired (TR/TE = 3390/389 ms; FOV = 256 mm; 1 mm³ isotropic).

#### Functional imaging

BOLD images for the fLoc and retinotopy runs were acquired with 1.5 mm3 isotropic voxel size and the following protocol parameters: TR/TE = 2000/36 ms, FA = 78°, FOV = 138 × 138 mm with 159 volumes per run (the first six were removed before analysis), and 80 slices. Because the acquisition did not cover the whole head, a saturation band was applied in the prefrontal region to prevent aliasing artifacts. Magnitude and phase images, as well as an additional volume for NORDIC denoising (Dowdle et al., 2021; Moeller et al., 2021; Dowdle et al., 2023), were acquired, and a single-band reference image was collected to improve registration to the structural scan. In addition to functional scans, each session included spin-echo EPI images acquired with opposite phase-encoding directions to estimate B0 field map and enable post hoc correction of susceptibility-induced distortions using a blip-up/blip-down approach (same overall slice slab as the fMRI GE-EPI data, 1.5 mm3 isotropic resolution, TR= 14926 ms, TE = 122.40 ms, flip angle 90°, refocusing flip angle 180° bandwidth 1132 Hz per pixel, partial Fourier 6/8, TA = 15 sec per direction).

For the preprocessing of the data, DICOMs were converted to BIDS (RRID:SCR_016124) using HeuDiConv 1.3.2 (RRID:SCR_017427). Functional MRI data were preprocessed using fMRIPrep 25.1.1 (RRID:SCR_016216), including T2-weighted images to improve FreeSurfer’s (RRID:SCR_001847) anatomical segmentation. For each subject, a single session’s T1-weighted scan was used as the anatomical reference across all sessions, ensuring that all functional data were registered to the same anatomical space. Surface-based time series were generated in each participant’s *fsnative* space using *bbregister* and all subsequent analyses were performed on the surface. No spatial smoothing was applied.

### MRI data analysis

#### GLM analysis and ROI delineation

First-level GLMs were estimated with NiLearn 0.10.4 (RRID:SCR_001362) and included confound regressors from fMRIPrep: 6 motion regressors, framewise displacement, unsteady-state volumes, and a/tCompCor components explaining 50% of variance (Figure 6, panel A). For delineating vOTC reading ROIs, we developed a new automated ROI delineation method. After GLM estimation, we combined five sessions from each subject and applied a watershed-based algorithm within a predefined vOTC search space.

**Figure 6.**
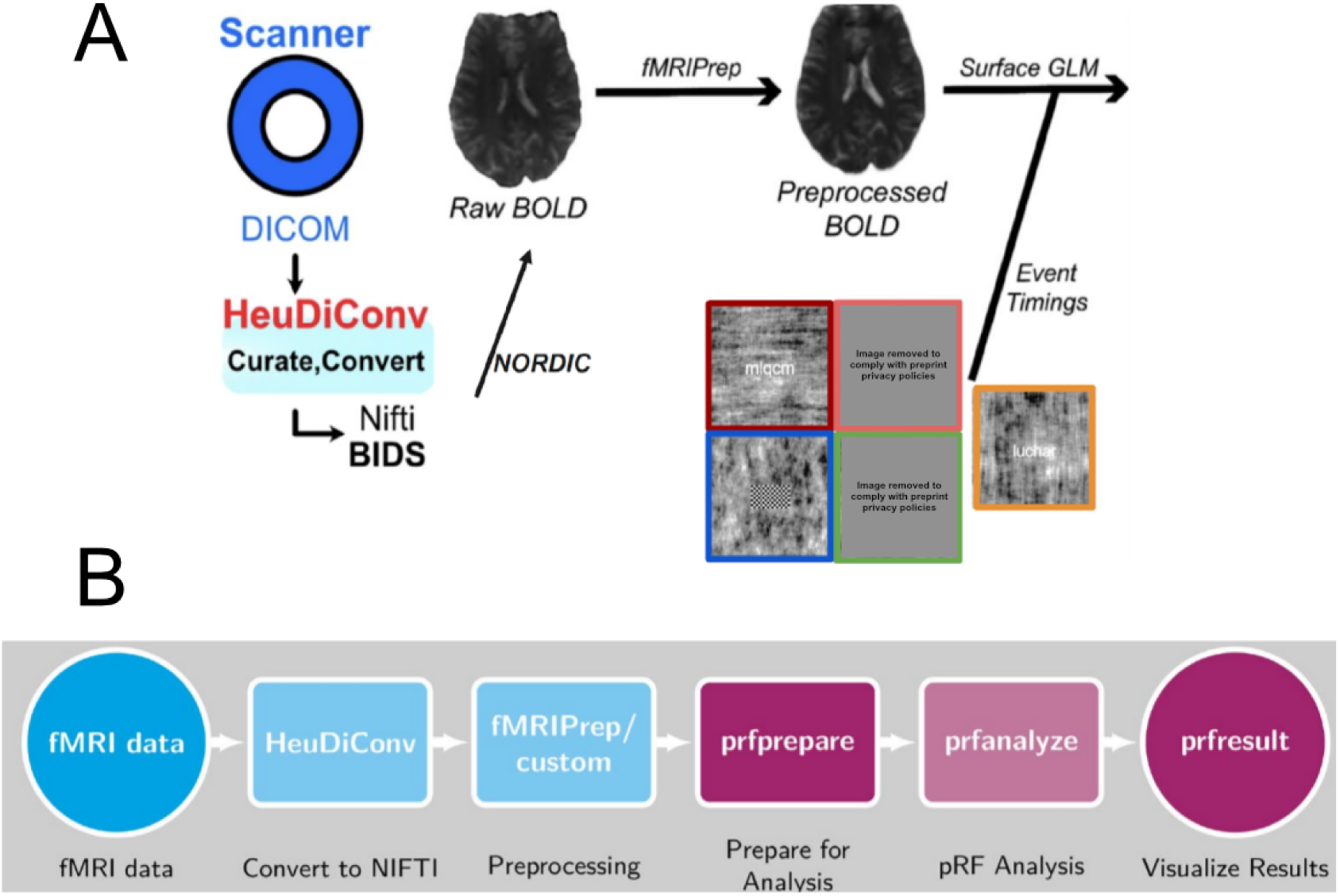
fMRI analysis pipeline. Common analysis stages: convert DICOMs to NIfTI and BIDS, denoising with NORDIC, preprocessing and transformation to the surface with fmriprep. **A.** fLoc analysis additionally performed GLM analysis using Nilearn. **B.** Retinotopy analysis additionally used *prfprepare*, *prfanalyze* and *prfresult* containers. *prfprepare* takes the preprocessed functional data in anatomical or surface space and prepares it for the pRF analysis. *prfanalyze* performs pRF analysis using vistasoft. *prfresult* plots the results as coverage maps or surface overlays. Reprinted from Linhardt et al. (2025).

Specifically, the T-values were scaled within this search space. This scaling step was necessary due to a pronounced anterior-to-posterior gradient in T-values, as well as high inter- and intra-subject variability, which precluded the use of a single, fixed T-value threshold across the entire sample or cortex. To implement this, we applied a local kernel during the scaling procedure, ensuring that every vertex was scaled relative to its closest neighbors. Following this local scaling, the standard deviation of the scaled T-value across sessions was calculated for each voxel. This yielded a voxelwise score that reflected both task-related activation strength and session-to-session variability. We then applied a reverse-watershed procedure to automatically identify candidate clusters.

Word-selective ROIs were manually defined on each subject’s native cortical surface (*fsnative*) in the left hemisphere by a trained rater (YL). We adopted a surface-based, single-subject approach because it preserves the fine-grained topographic organization of the vOTC and aligns with anatomical landmarks previously shown to separate adjacent functional subregions. Based on prior studies of the vOTC, five candidate word-selective ROIs were delineated per subject and organized along the posterior-anterior axis: OWA, pOTS, mOTS, aOTS and mFus. The full set of ROIs, together with their anatomical landmarks and defining contrasts, is illustrated in Figure 7.

**Figure 7.**
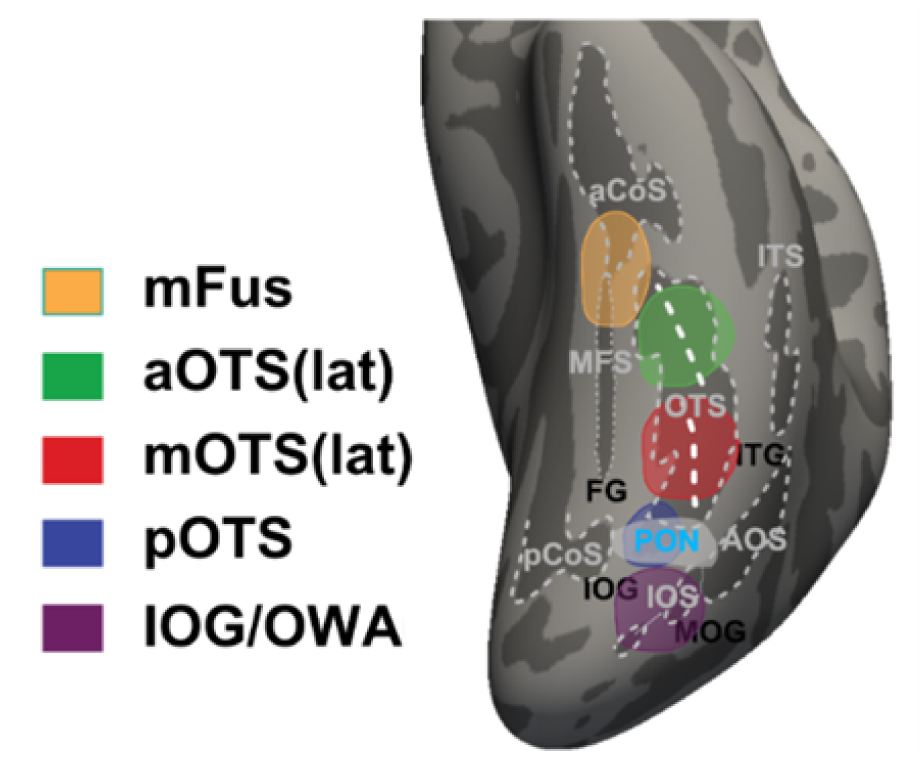
vOTRC ROIs and Cross-Subject Variable Topology. Template brain with the 5 functional regions of interest (ROIs) consistently identified across all 11 subjects in native space. Note that the a- and m-OTS can be reliably segmented into distinct lateral subregions (lat-aOTS and lat-mOTS) in only a subset of the cohort (∼50% of subjects). Reprinted from Lerma-Usabiaga et al. (2026).

ROIs were labeled using a custom HTML tool based on auto clusters using score maps derived from three functional contrasts: real words versus scrambled words (RW > SC), real words versus false-fonts (RW > FF), and faces versus all other categories (Faces > others). Score maps were obtained from subject-level t-maps using a rescaling-and-scoring procedure described in Lerma-Usabiaga et al. (2026).

Labelling proceeded in posterior-to-anterior order along the occipito-temporal sulcus (OTS) and the mid-fusiform sulcus (MFS); ROI boundaries were guided by local functional clusters, sulcal landmarks, the Destrieux cortical parcellation (Destrieux et al., 2010), and the spatial relationship between word-selective and face-selective regions. Three face-selective ROIs were first identified using the (Faces > others) contrast. The occipital face area (OFA) was defined as the most posterior face-selective cluster in the inferior occipital gyrus. FFA-1 was defined as the face-selective cluster near the posterior tip of the MFS, whereas FFA-2 was defined as the cluster near the anterior tip of the MFS. These landmarks were stable across subjects and are consistent with previous anatomical and functional descriptions of ventral temporal face-selective cortex (Chen et al., 2023; Weiner et al., 2014).

Three word-selective ROIs were then delineated relative to these face-selective anchors. mFus, mOTS, and OWA were defined as (RW > SC) word-selective clusters located near FFA-2, FFA-1, and OFA, respectively. This spatial arrangement was consistent across the 11 subjects. The close proximity between FFA-2 and mOTS is also consistent with previous descriptions of the VWFA and adjacent face-selective cortex (Dehaene & Cohen, 2007).

aOTS was defined as the word-selective cluster located between mFus and mOTS along the anterior-posterior axis, near the anterior portion of the OTS. pOTS was identified as the cluster showing positive responses in the (RW > SC) contrast but showing no responses or negative responses in the (RW > FF) contrast. This two-contrast intersection criterion follows Lerma-Usabiaga et al. (2018) and isolates the VWFA subregion associated with visual-feature extraction. However, the separability of the anterior and middle OTS clusters varied across subjects. aOTS could be separated from neighboring word-selective cortex in 6 of 11 subjects, whereas mOTS showed a separable pattern in 5 of 11 subjects. Because of this variability, we treated the distinction between a five-ROI and seven-ROI scheme as uncertain during ROI definition and in this work we only used the 5 reading ROIs version.

#### Retinotopy analyses

Following preprocessing, the surface time series were prepared for pRF analysis (Figure 6, panel B) using the *prfprepare* container (Lerma-Usabiaga et al., 2020). This step converted the stimulus presentation logs into NIfTI-format stimulus apertures and masked each subject’s surface data according to the Wang et al. (2015) probabilistic atlas, restricting subsequent analyses to the 22 ROIs of interest (V1v, V2v, V3v, hV4, Vo-1, V1d, V2d, V3d, V3a, IPS-0, IPS-1; all with their left and right hemisphere regions). Masked time series for each subject, session, and hemisphere were saved as individual files, together with a sidecar record specifying each ROI’s vertex indices, allowing results to be reconstructed and grouped by region at later stages of the pipeline.

pRF parameters were then estimated using the *prfanalyze* container (Lerma-Usabiaga et al., 2020), which implements the vistasoft fitting algorithm. For each vertex, the model’s center coordinates (x, y) and receptive field size (σ) were estimated through a coarse-to-fine fitting procedure, using the One Gaussian (OG) model (Dumoulin & Wandell, 2008) as the primary framework. A parallel analysis using the Compressive Spatial Summation (CSS) model (Kay et al., 2013) was conducted for later model result comparisons. Eccentricity was computed for each vertex as the Euclidean distance between its estimated pRF center and the point of fixation. Resulting pRF parameter maps were reconstructed and visualized using the *prfresult* container.

#### Word-Center stimulus analyses

Data from sessions that include the Word-Center paradigm were analyzed simultaneously with the two previous analysis pipelines: (1) the same GLM approach used for the fLoc runs to extract localizer-style contrasts and to identify reading-responsive ROIs (Figure 6, panel A), and (2) the same pRF pipeline used for the dedicated retinotopy runs, yielding pRF parameter estimates from the hybrid Word-Center stimuli (Figure 6, panel B). In this case, the pRF estimates were obtained in both the 22 ROIs (Wang et al., 2015) previously described and in the 5 reading ROIs (Figure 7).

By processing the Word-Center data through both analysis frameworks (Nilearn GLMs and mrVista pRF modeling) we obtained two complementary descriptions of the same data: one emphasizing condition contrasts and localizer-defined ROIs, and the other providing retinotopic parameters.

### Comparative Statistical Analyses

#### Retinotopic comparison filtering parameter selection

To ensure reliable pRF estimates and maintain consistency across stimulus conditions, vertices were filtered based on model fit quality and spatial plausibility prior to analysis. The following thresholds were applied:

#### Goodness-of-Fit (R^2^)

Only vertices where the pRF model explained at least 20% of the variance in the BOLD time series were included, matching the threshold used in the precedent study of these analyses (Lerma-Usabiaga et al., 2021). This variance-explained criterion varies across studies (Benson et al., 2018; Gomez et al., 2018). The selected threshold balances rigorous reliability with the need for an adequate sample size across the cortical hierarchy, particularly in early visual areas and anterior ventral regions where signal-to-noise ratios may vary.

#### pRF Size

The Gaussian spread parameter (σ) was restricted to 1–15 degrees, in line with the range of biologically plausible pRF sizes reported across the visual hierarchy (Dumoulin & Wandell, 2008; Harvey & Dumoulin, 2011). This range excludes overfitting artifacts from implausibly small pRFs and non-specific responses from excessively large pRFs that may span multiple visual field representations.

#### Eccentricity

Inclusion was limited to vertices with eccentricity estimates between 4 and 15 degrees, matching the extent of the mapping stimulus used in this and prior studies of the reading circuitry (Le et al., 2017; Lerma-Usabiaga et al., 2021). Vertices below 4 degrees were excluded to avoid foveal saturation effects and the high representation density at the fixation point, while those above 15 degrees were removed to avoid edge effects outside the stimulus window.

To perform meaningful comparisons between different stimulus conditions, we implemented a vertex-pairing logic per ROI. For any pairwise analysis, a vertex was only included if it independently passed the three quality filters (R^2^, σ, and eccentricity) in both compared conditions. This strict pairing ensures that mean eccentricity bias calculations are performed on an identical neural population across conditions, preventing results from being driven by the selection of different vertices. This approach provides a conservative and robust measure of how the same cortical site reconfigures its spatial tuning in response to varying sensory and cognitive demands.

### Retinotopic comparison metric selection

The most direct way to visualize pRF eccentricity differences between conditions is through per-vertex-pair scatterplots (Figure 3), where each point represents a single cortical vertex plotted according to its eccentricity estimate in each of the two conditions (one per axis), so that deviations from the diagonal reflect differences in spatial tuning between conditions. However, given the scale of the present dataset (spanning 22 ROIs, multiple FWS configurations, stimulus comparisons, and 11 participants), scatterplots alone would produce an unmanageable number of panels. As our central question was whether one stimulus condition systematically shifted pRF estimates toward the fovea or periphery relative to another, we selected a metric directly summarizing that average directional shift: the mean bias in degrees of visual angle. For each ROI and condition pair, this was defined as the mean signed difference in per-vertex eccentricity estimates. Importantly, ROIs with fewer than 25 vertex-pairs were excluded from the analysis to ensure that the mean eccentricity bias calculated for each region was statistically robust and representative of the underlying neural population. Statistical significance was assessed using a linear mixed-effects model with a subject random intercept to account for the repeated-measures structure of the data and a pair variance component to account for variability between session pairs. This allowed the full pattern of results to be presented compactly in color-coded matrices, with rows indexing experimental conditions and columns indexing ROIs.

### Classical retinotopy analyses in visual and reading ROIs

The analyses described in this section were applied to data from classical retinotopic mapping sessions, in which a flickering ring aperture traversed the visual field using one of three carriers: CB, RW, or FF. Each session was additionally characterized by an FWS parameter triplet that specified the temporal frequency, bar width, and element size of the mapping aperture, allowing us to examine how these low-level stimulus properties influence eccentricity estimates independently of carrier type.

#### FWS-level stimulus carrier comparison analysis

To quantify stimulus-dependent variations in spatial tuning, we compared the pRF eccentricity estimates obtained with the three carrier types (CB, RW, and FF) across all ROIs. Scanning sessions were paired according to identical FWS configurations, so that any differences in eccentricity between conditions reflected the carrier type rather than the spatial properties of the mapping aperture. Mean eccentricity bias was then computed for each pairwise stimulus comparison (CB–RW, CB–FF, and FF–RW) in each ROI, allowing us to track how stimulus sensitivity changes along the visual hierarchy. A primary goal of this analysis was to replicate the hierarchical stimulus-dependency pattern reported by Lerma-Usabiaga et al. (2021), in which RW stimuli consistently produced more foveal pRF center estimates than CB (CB ecc > RW ecc), with the divergence growing progressively larger along the ventral visual pathway (from V1v to VO-1).

#### Subject-level stimulus carrier comparison analysis

Given the known inter-individual variability in pRF spatial preferences (Glezer & Riesenhuber, 2013), we additionally conducted the stimulus comparison analysis at the single-subject level. Within each participant, all available classical retinotopic sessions were paired regardless of their FWS configuration, maximizing the number of within-subject comparisons. This approach allowed us to assess the consistency of stimulus-dependent eccentricity shifts across individuals and to quantify the proportion of participants showing the expected hierarchical pattern. As a consequence of pooling across FWS configurations, within-subject variability in these comparisons reflects not only carrier-type effects but also the influence of differences in temporal frequency and element size between paired sessions.

In addition to the comparison matrices described above, we generated a complementary set of subject-level plots to perform the carrier-type comparisons in the 5 vOTRC reading ROIs (Figure 7), alongside the left ventral visual pathway (V1v to VO-1). The first set of plots examined the three classical retinotopic carrier-type contrasts (CB–RW, CB–FF, FF–RW), plotting each subject’s mean eccentricity bias, rather than a single aggregated value, at each ROI. This allows the magnitude and direction of a given carrier-type bias to be tracked continuously from the ventral visual pathway into the reading circuitry.

The second set of plots used the same format to examine each carrier-type (RW, FF, CB) individually rather than as pairwise contrasts, plotting each subject’s mean eccentricity within a single stimulus type at each ROI. Because this analysis does not involve a between-condition comparison, no session-pairing was required: all available sessions of the relevant carrier-type were used for each subject regardless of FWS configuration. This allowed us to examine whether the absolute eccentricity tuning of each carrier-type, and not only the bias between pairs of carrier-types, follows a systematic gradient when moving from the ventral visual pathway into the reading circuitry.

Because the reading ROIs, and the more anterior vOTRC regions in particular, retain comparatively few vertices under our standard filtering criteria, leaving several subjects and ROIs sparsely represented or untestable, we relaxed the goodness-of-fit and eccentricity thresholds for this specific set of plots (R² > 0.05; minimum eccentricity > 1°), with all other criteria unchanged. This relaxed filtering was applied only to the gradient plots described here and should be treated as exploratory.

#### Stimulus frequency and size effect analysis

As a secondary, exploratory analysis, we examined whether the flickering frequency and element size components of the FWS parameter independently influenced eccentricity estimates. Pairwise session comparisons were constructed by holding one parameter constant while varying the other (e.g., comparing 2 Hz against 8 Hz at a fixed size). Mean eccentricity bias was calculated for each contrast. Because the number of available sessions was unequal across FWS combinations, these comparisons were treated as exploratory and interpreted with appropriate caution.

#### pRF eccentricity reliability

The stability of pRF eccentricity estimates was assessed using Lin’s CCC (Lin, 1989) applied to vertex-level eccentricity estimates. For each FWS configuration, we identified all subjects contributing at least two sessions and formed all unique within-subject session-pairs. For each (subject, session-pair, ROI, stimulus) cell, vertices present in both sessions were matched by index, retaining only vertices that passed the previously described filtering criteria in both sessions.

Lin’s CCC was then computed on the paired eccentricity estimates (session A vs. session B) for each retained cell, yielding one CCC value per (subject, session-pair, ROI, stimulus) combination. These cell-level values were aggregated into a single summary value per FWS configuration through a hierarchical cascade of medians: across session-pairs within each (subject, ROI, stimulus) combination, then across stimuli within each (subject, ROI), then across subjects within each ROI, and finally across ROIs, yielding one pooled CCC per FWS configuration. Consistent with eccentricity mean bias analyses, session-pairs with fewer than 25 vertex-pairs were excluded.

The traditional Intraclass Correlation Coefficient (ICC) was not used to assess the reliability because our data structure involves multiple stimulus types, ROIs, and session types (FWS) with unbalanced observations. Despite not being perfect, the CCC approach maximizes statistical power by allowing us to pool reliability estimates across all available session pairs using median aggregation. In contrast, ICC requires balanced designs limited to within-subject comparisons or complex mixed-effects modeling to handle heterogeneous variance structures across conditions, which would reduce the effective sample size and complicate interpretation in our multi-factor design.

Two FWS configurations (4|3|25 and 4|3|100) were excluded from the reliability analysis, as no subject contributed two or more sessions under either configuration, leaving no within-subject session-pairs to compare. For the remaining seven classical retinotopy conditions, plus the WC (8|3|50) condition, 95% bootstrap confidence intervals (2000 replicates) were obtained by randomly resampling subjects with replacement and recomputing the full CCC cascade on each replicate, providing a non-parametric estimate of uncertainty around each pooled CCC point estimate.

#### pRF model comparison: OG vs CSS

Although the One Gaussian (OG) model was used as the primary estimation framework (Dumoulin & Wandell, 2008), we performed a parallel analysis using the Compressive Spatial Summation (CSS) model (Kay et al., 2013) to verify if the choice of framework influenced our spatial tuning estimates. The CSS model extends the OG approach by incorporating a compressive nonlinear response exponent that captures subadditive spatial summation, a property that becomes increasingly prominent in higher-level visual areas (Le et al., 2017). Mean eccentricity bias between the two models was computed across all stimulus types, FWS configurations and all 22 ROIs.

### Word-Center paradigm analyses

The Word-Center (WC) paradigm was designed to simultaneously engage bottom-up sensory and top-down linguistic processing. pRF estimates from these sessions were analyzed using the same mean eccentricity bias metric applied to the classical retinotopy data.

#### WC stimulus carrier comparison in visual and reading regions

To characterize how the nature of the fixated stimulus affects spatial tuning during simultaneous sensory and linguistic stimulation, we compared pRF eccentricity estimates across the three WC conditions (fixRW, fixFF, and fixRWblock). Mean eccentricity bias was computed for each pairwise comparison across all ROIs. This analysis aimed to reveal whether the linguistic content of the attended stimulus or its spatial arrangement (continuous string vs. blocked) systematically shifts pRF center estimates.

As with the classical retinotopy analyses above, we also examined these three WC contrasts, as well as each WC condition individually, using the subject-level gradient plots described in the “Subject-level stimulus carrier comparison analysis” section, restricted to the vOTRC reading-related ROIs and the left ventral visual pathway

#### Cross-paradigm comparisons: classical retinotopy vs. Word-Center in visual and reading regions

To assess how the concurrent presentation of sensory and linguistic information modifies pRF estimates relative to classical retinotopic mapping, we compared eccentricity values derived from the two paradigms, pairing only the frequency and size equivalent 8|3|50 classical retinotopy and WC (8|3|50) sessions within each subject (Figure 5). To ensure stimulus compatibility, comparisons were restricted to pairs sharing at least one component: CB against all three WC conditions, RW against fixRW and fixRWblock, and FF against fixFF. Mean eccentricity bias was computed for each cross-paradigm pairing across all ROIs.

As above, we also examined these cross-paradigm contrasts using the same subject-level gradient plots described in the “Subject-level stimulus carrier comparison analysis” section, restricted to the vOTRC reading ROIs and the left ventral visual pathway.

#### WC localizer values in reading regions

To validate whether the WC fixRWblock condition is a suitable tool for identifying specialized reading areas, we performed a functional overlap analysis using the 5 reading-related ROIs obtained with the fLOC localizer (OWA, pOTS, mOTS, aOTS, and mFus; Figure 7). This analysis was restricted to these reading-selective ROIs and was not extended to the retinotopic visual ROIs (Wang et al., 2015), as the WC paradigm is designed specifically to engage reading-related processing, making a comparable validation within purely visual regions uninformative. For each subject, we computed a real-word-versus-null (RWvsNull) contrast restricted to the central fixation content of the fixRWblock condition: the blocks in which real words were presented at fixation were contrasted against the alternating blank blocks in which no word was shown at fixation. Because the peripheral moving checkerboard bar was present continuously throughout the run, independent of this alternation, it was common to both block types and therefore did not contribute to the contrast, which isolates the response to the presence versus absence of central linguistic content. This yielded a mean T-value per ROI that reflects the response evoked by the word blocks relative to baseline. The T-values indicate whether the spatial tuning estimates derived from the Word-Center paradigm are rooted in the same neural populations that define the core reading network. This analysis was restricted to session 10, the only session common to all 11 participants, and to the left hemisphere, where the reading circuitry is most consistently lateralized. Group-level results were visualized as the across-subject distribution of mean T-values within each ROI, together with individual subject values and the group median.

Although data was also collected for the continuous fixRW condition, the fixRWblock condition was selected for this analysis as it provided a higher signal-to-noise ratio. The fixRW results exhibited significantly more noise, likely due to neural habituation resulting from the constant presentation of orthographic material.

## Results

### Classical retinotopy analyses in visual and reading regions

#### FWS-level stimulus carrier comparison analysis

To examine how carrier-types affect spatial tuning across the visual and reading-related hierarchy, we compared pRF eccentricity estimates for CB, RW, and FF stimuli of subjects grouped by FWS configurations.

In the visual regions, differences in pRF eccentricity across carrier-types tended to increase along the visual hierarchy, with the largest biases observed in higher-order areas (hV4, VO-1, IPS-0, IPS-1) in both ventral and dorsal streams (Figure 8). Early areas (V1–V3) showed comparatively smaller deviations. The effect was driven primarily by differences between orthographic stimuli (RW and FF) and checkerboards (Figure 8, panels A and B); comparisons between RW and FF (Figure 8, panel C) revealed minimal differences.

**Figure 8.**
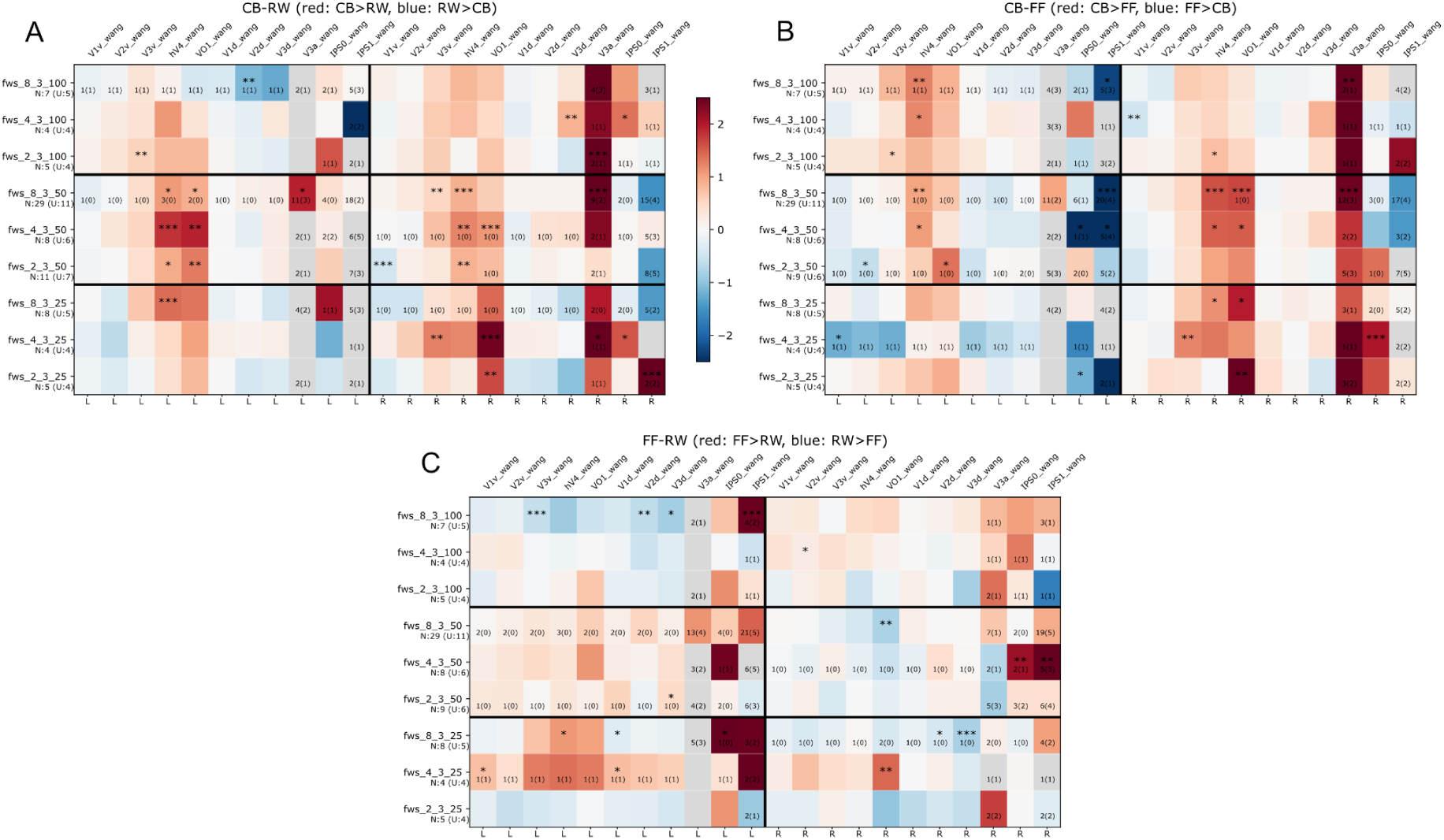
Stimulus carrier comparison across all FWS configurations and visual ROIs. **A.** CB−RW. **B.** CB−FF. **C.** FF−RW. Rows represent the nine FWS conditions, grouped by size block (largest to smallest, top to bottom), with bold horizontal lines separating blocks and frequency varying within each block. Columns represent the 22 ROIs from the Wang atlas; left-hemisphere ROIs occupy the first 11 columns and right-hemisphere ROIs the next 11, separated by a bold vertical line. Within each hemisphere, ventral-stream areas (V1v-VO1) precede dorsal-stream areas (V1d-IPS1), placing higher-order areas toward the right of each cluster. Cell color follows a diverging red/blue scale (±2.5°), showing mean eccentricity bias (Y − X, degrees of visual angle) between the paired stimuli types: red indicates Y > X (more peripheral eccentricity for the first stimulus), blue indicates X > Y, and gray indicates no testable vertices. Significance was assessed per cell using a linear mixed-effects model of per-session-pair differences with a subject random intercept; stars mark cells where the null hypothesis of zero bias is rejected at p < 0.05 (***p < 0.001, **p < 0.01, *p < 0.05). Cells with fewer than two paired sessions or less than 25 paired-vertices were not tested. Row labels (N:x, U:y) indicate the number of contributing sessions (N) and unique subjects (U) for each FWS condition; per-cell deficit markers in the lower-right corner flag individual ROIs where the local sample falls below the row N(U) baseline.

Across the ventral hierarchy (V1–VO-1), pRF centers for CB were systematically more peripheral than those for RW (Figure 8, panel A) in 7 out of 9 FWS types (77%), with this divergence present bilaterally and becoming increasingly pronounced in higher-order maps.

#### Subject-level stimulus carrier comparison analysis

To assess the consistency of these carrier-type effects across individuals, we examined stimulus-dependent eccentricity shifts at the single-subject level.

In the visual regions, 7 out of 10 participants (70%; subjects 2, 3, 4, 6, 7, 8, and 9) showed a systematic foveal shift for RW stimuli compared to CB across the ventral pathway of both hemispheres (Figure 9, panel A). Comparisons between RW and FF (Figure 9, panel C) revealed minimal differences. Subject 11 was excluded from this count, as they contributed only a single scanning session (Figure 5); nevertheless, the expected hierarchical gradient was present in this subject’s left hemisphere. It should be noted that, unlike the FWS-level analysis in the previous section, these per-subject results aggregate sessions across different FWS configurations, meaning that variability in temporal frequency and element size may contribute to within-subject variance.

**Figure 9.**
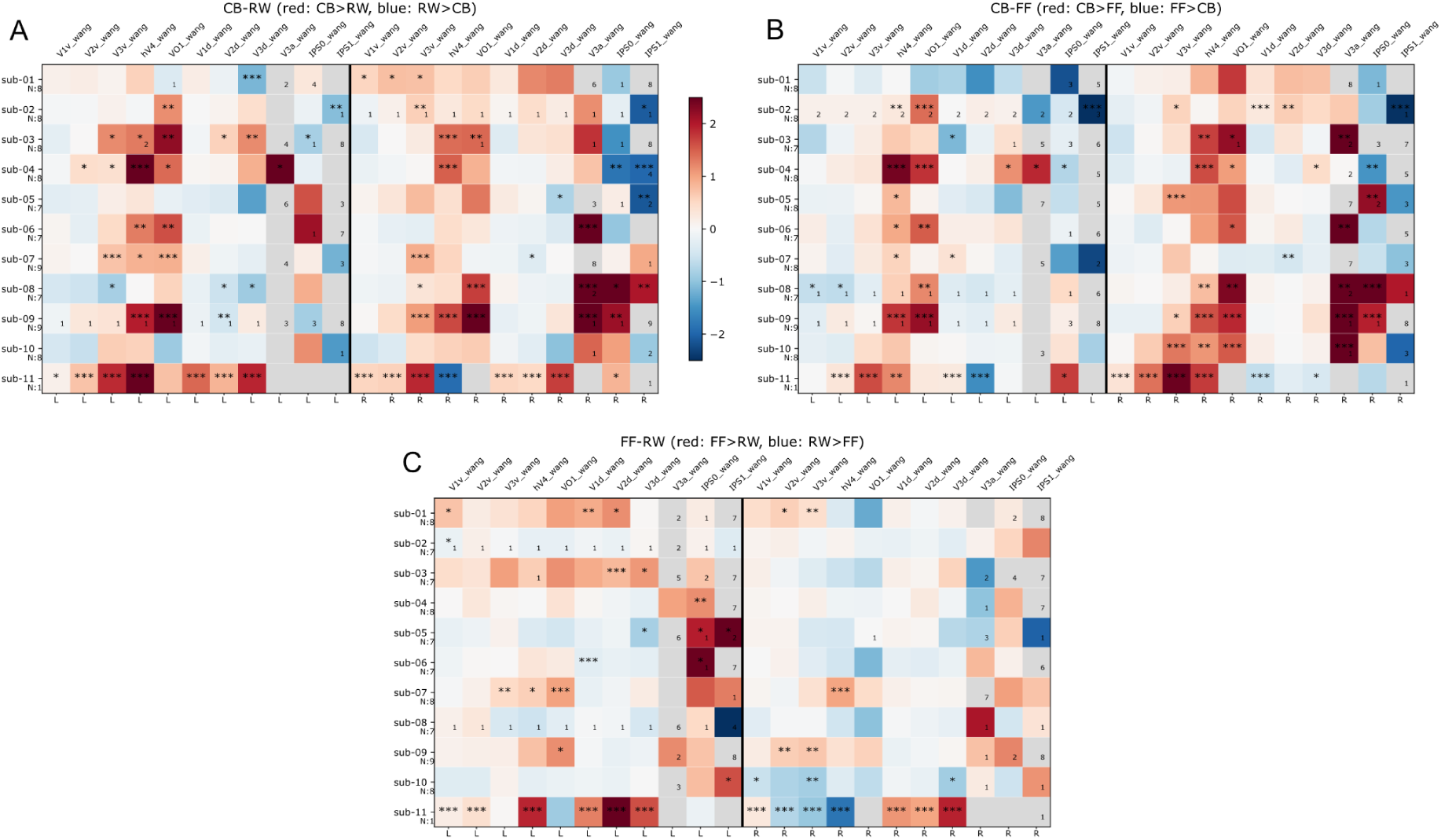
Stimulus carrier comparison across all subjects and visual ROIs. **A.** CB−RW. **B.** CB−FF. **C.** FF−RW. Rows represent the 11 subjects in numerical order. Columns represent the 22 ROIs from the Wang atlas; left-hemisphere ROIs occupy the first 11 columns and right-hemisphere ROIs the next 11, separated by a bold vertical line. Within each hemisphere, ventral-stream areas (V1v–VO1) precede dorsal-stream areas (V1d–IPS1), placing higher-order areas toward the right of each cluster. Cell color follows a diverging red/blue scale (±2.5°), showing mean eccentricity bias (Y − X, degrees of visual angle) between the paired stimuli types: red indicates Y > X (more peripheral eccentricity for the first stimulus), blue indicates X > Y, and gray indicates no testable vertices. Significance was assessed per cell using a linear mixed-effects model of per-session-pair differences (as sub-11 only has one session, the mean bias is calculated without the mixed effects model and the significance comes from a t-test); stars mark cells where the null hypothesis of zero bias is rejected at p < 0.05 (***p < 0.001, **p < 0.01, *p < 0.05). Cells with fewer than two paired sessions or less than 25 paired-vertices were not tested. Row labels (N) indicate the number of contributing sessions for that subject; per-cell deficit markers in the lower-right corner flag individual ROIs where the session sample falls below the row N baseline.

Across both the ventral visual pathway and the reading-specific regions, the pattern of eccentricity estimates varied systematically depending on the carrier type (Figure 10). When examining the carrier mean eccentricity values individually (Figure 10, panel A), we observe a consistent pattern across ROIs and carriers: a continuous decrease in eccentricity from V1v to hV4, followed by a spike in VO-1. In the reading ROIs, eccentricity starts at very low levels in the OWA and increases as one moves anteriorly through the hierarchy. The only exception is mFus for RW and FF stimuli, where eccentricity is not higher than in the preceding aOTS. Additionally, mFus is generally situated more medially and slightly more anteriorly than the aOTS.

**Figure 10.**
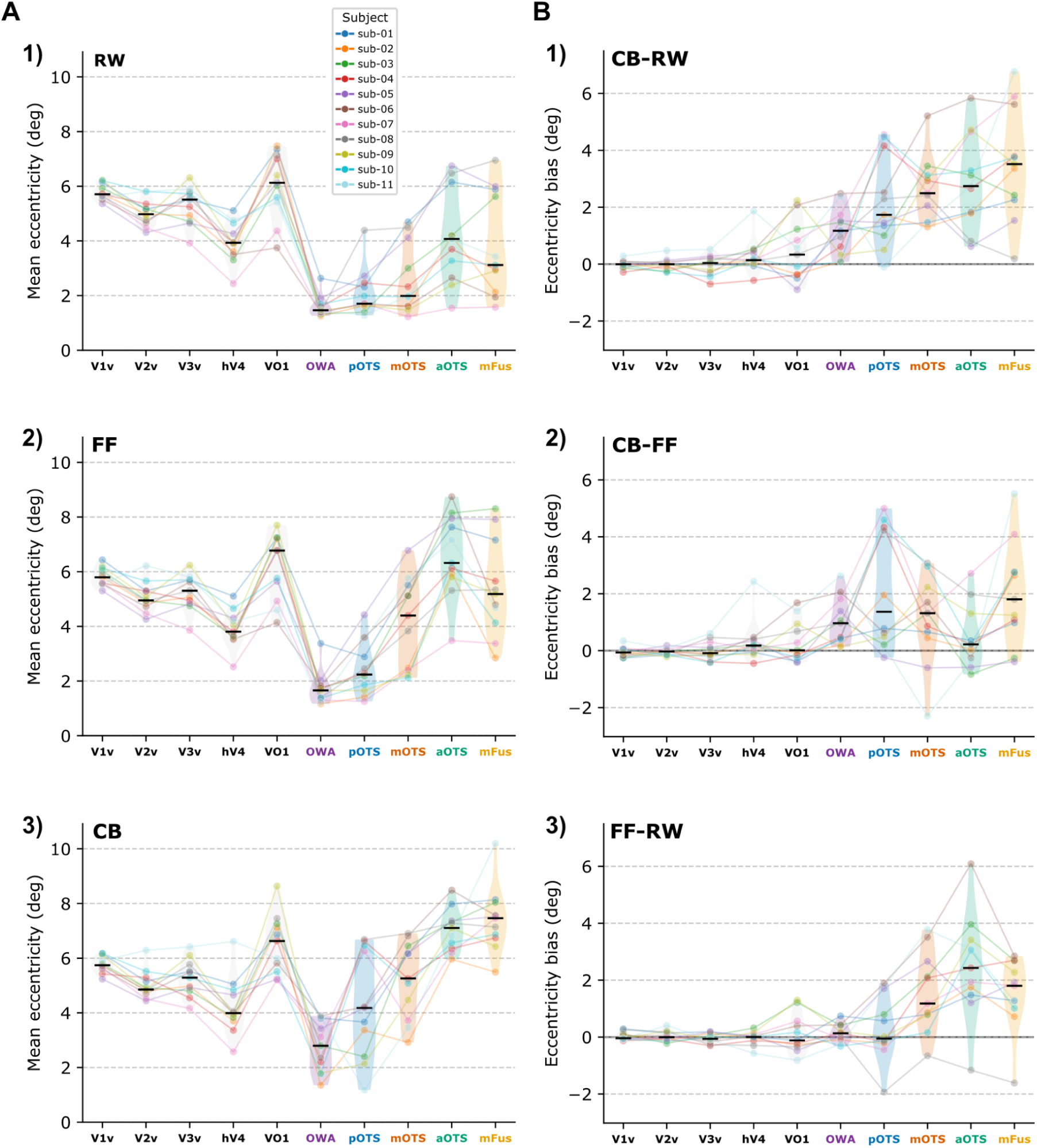
Classical retinotopy stimulus carrier comparison across all subjects and ventral visual and reading ROIs. Individual colored dots and connecting lines represent the 11 participants; black lines indicate the group median within each ROI. The X-axis tracks 10 ventral ROIs ordered from posterior to anterior. **A.** Y-axis shows the mean eccentricity per subject (in degrees). Rows display results for different carriers within the stimulus bar: 1) RW, 2) FF, and 3) CB. **B.** Y-axis shows the subject-specific eccentricity bias between carriers (in degrees). Rows display carrier differences: 1) CB−RW, 2) CB−FF, and 3) FF−RW.

Comparing carriers based solely on the eccentricities of individual ROIs is difficult. Therefore, in Fig. 10, panel B, we plot the pairwise carrier eccentricity comparisons. While all comparisons followed a posterior-to-anterior gradient, they exhibited distinct behaviors. The CB vs. RW comparison shows a clear gradient, progressing from early visual fields, where differences are minimal, to sustained increases throughout the reading regions of interest; in all instances, RW estimates are more foveal than CB. The comparison between FF and CB follows a similar pattern with two notable exceptions: (1) the magnitude of the difference is smaller, and (2) the aOTS shows practically no difference. Finally, the FF vs. RW comparison shows no significant differences in visual areas or posterior reading regions (OWA, pOTS), but reveals an approximate 2-degree difference in the three anterior regions (mOTS, aOTS, and mFus).

#### Stimulus frequency and size effect analysis

To examine whether the temporal and spatial properties of the mapping aperture also contribute to eccentricity estimates, we compared sessions differing only in temporal frequency or element size while holding the remaining parameters constant.

Lower flickering frequencies tended to yield more peripheral eccentricity estimates. At sizes 25 and 50 (Fig. 11, panels A and B), lower temporal frequencies (2 Hz) consistently produced greater eccentricity than higher ones (8 Hz), with 2–8 Hz and 4–8 Hz contrasts being most robust. This effect was absent at size 100 (Fig. 11, panel C), suggesting a frequency by size interaction that requires further investigation.

**Figure 11.**
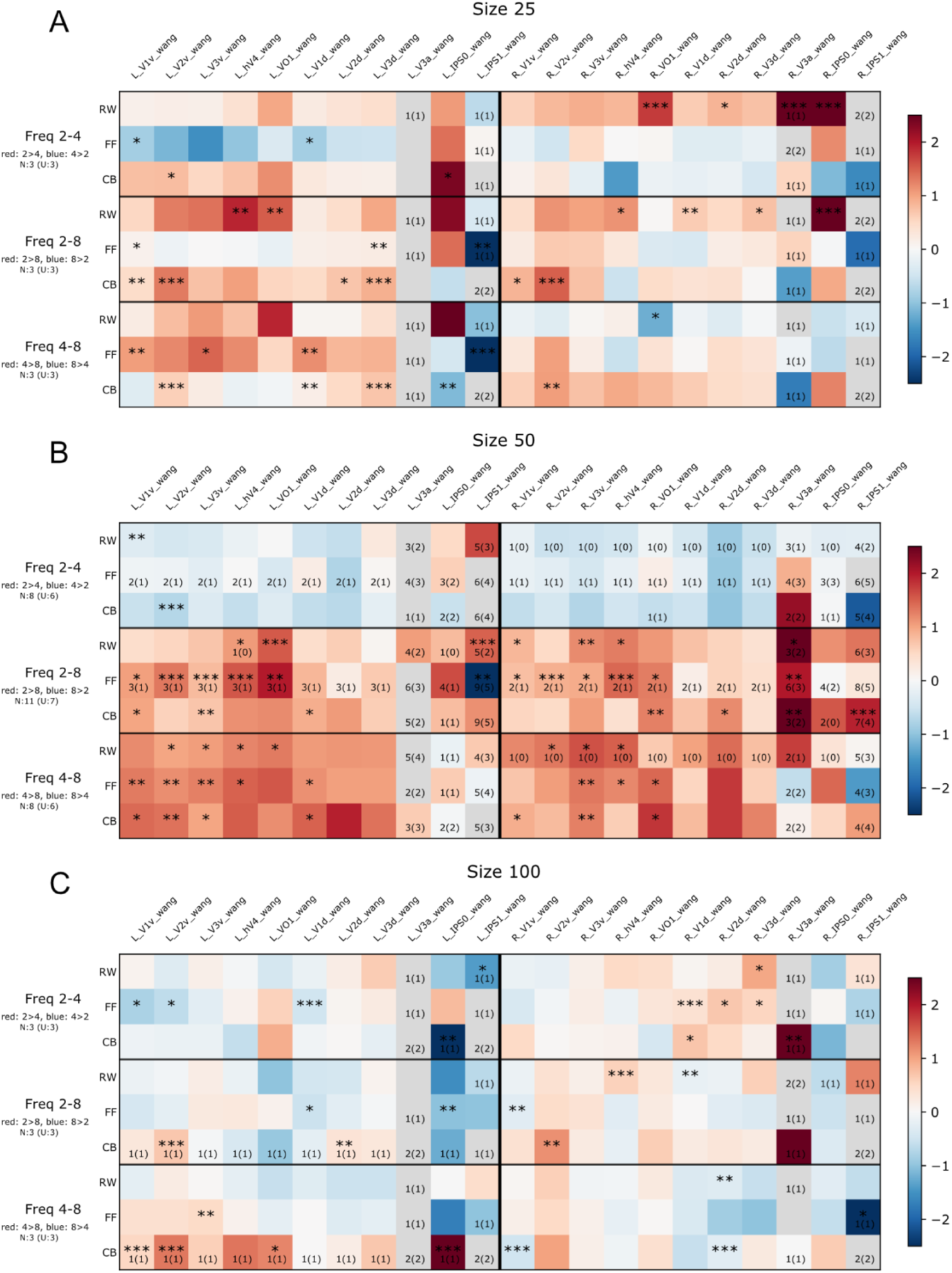
Frequency effect comparison matrices across all sizes and visual ROIs. **A.** Size 25. **B.** Size 50. **C.** Size 100. Row triplets represent the three frequency pair comparisons (2-4, 2-8, 4-8), with one of the three stimulus types per row (RW, FF, CB). Columns represent the 22 ROIs from the Wang atlas; left-hemisphere ROIs occupy the first 11 columns and right-hemisphere ROIs the next 11, separated by a bold vertical line. Within each hemisphere, ventral-stream areas (V1v–VO1) precede dorsal-stream areas (V1d–IPS1), placing higher-order areas toward the right of each cluster. Cell color follows a diverging red/blue scale (±2.5°), showing mean eccentricity bias (Y − X, degrees of visual angle) between the paired frequency values: red indicates Y > X (more peripheral eccentricity for the first frequency value), blue indicates X > Y, and gray indicates no testable vertices. Significance was assessed per cell using a linear mixed-effects model of per-session-pair differences with a subject random intercept; stars mark cells where the null hypothesis of zero bias is rejected at p < 0.05 (***p < 0.001, **p < 0.01, *p < 0.05). Cells with fewer than two paired sessions or less than 25 paired-vertices were not tested. Row labels (N:x, U:y) indicate the number of contributing sessions (N) and unique subjects (U) for each frequency pair; per-cell deficit markers in the lower-right corner flag individual ROIs where the local sample falls below the row N(U) baseline.

Smaller stimulus elements tended to produce more peripheral eccentricity estimates, though this effect was neither as strong nor as uniform as the frequency effect described above. The pattern was clearest for the largest size contrasts (25–50 and 25–100) at lower temporal frequencies (2 and 4 Hz), particularly along the ventral pathway (Fig. 12, panels A and B), while the 50–100 contrast and the 8 Hz frequency condition (Fig. 12, panel C) showed weaker and less consistent biases, at times reversing direction. Given the unequal number of sessions available across FWS combinations, the apparent hemispheric differences observed across both frequency and size conditions should be treated as exploratory observations requiring replication rather than robust findings.

**Figure 12.**
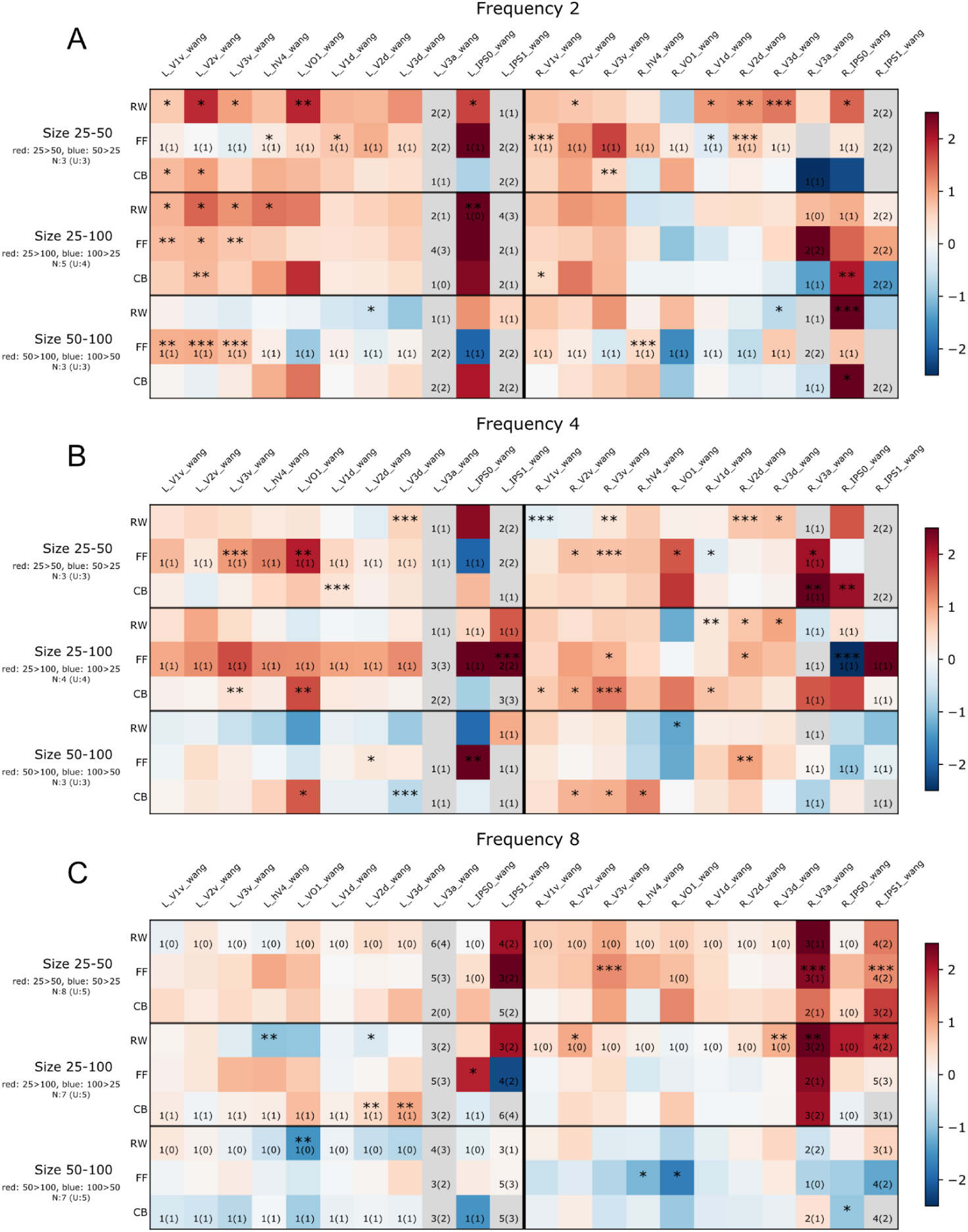
Size eccentricity comparison matrices across all frequencies and visual ROIs. **A.** Frequency 2. **B.** Frequency 4. **C.** Frequency 8. Row triplets represent the three size pair comparisons (25-50, 25-100, 50-100), with one of the three stimulus types per row (RW, FF, CB). Columns represent the 22 ROIs from the Wang atlas; left-hemisphere ROIs occupy the first 11 columns and right-hemisphere ROIs the next 11, separated by a bold vertical line. Within each hemisphere, ventral-stream areas (V1v–VO1) precede dorsal-stream areas (V1d–IPS1), placing higher-order areas toward the right of each cluster. Cell color follows a diverging red/blue scale (±2.5°), showing mean eccentricity bias (Y − X, degrees of visual angle) between the paired size values: red indicates Y > X (more peripheral eccentricity for the first size value), blue indicates X > Y, and gray indicates no testable vertices. Significance was assessed per cell using a linear mixed-effects model of per-session-pair differences with a subject random intercept; stars mark cells where the null hypothesis of zero bias is rejected at p < 0.05 (***p < 0.001, **p < 0.01, *p < 0.05). Cells with fewer than two paired sessions or less than 25 paired-vertices were not tested. Row labels (N:x, U:y) indicate the number of contributing sessions (N) and unique subjects (U) for each frequency pair; per-cell deficit markers in the lower-right corner flag individual ROIs where the local sample falls below the row N(U) baseline.

#### pRF eccentricity reliability

To confirm that the cross-session pairings used in the stimulus comparison analyses reflect genuine carrier-type effects rather than intersession measurement variability, we assessed the reproducibility of pRF eccentricity estimates across sessions sharing the same stimulus type and FWS configuration.

Test-retest reliability, assessed using Lin’s CCC on paired vertex-level eccentricity estimates, varied moderately across the eight tested conditions, with pooled CCC values ranging from 0.55 (8|3|100) to 0.71 (4|3|50), with varying confidence interval widths (Table 1). None of the conditions reached the high end of the CCC scale, suggesting that session-to-session agreement in pRF eccentricity estimates, while consistently positive, is best characterized as moderate rather than excellent across the mapping paradigms tested here.

**Table. 1.** pRF eccentricity reliability analysis. Condition: FWS configurations of the compared sessions; U: number of unique subjects included; N: number of sessions included; Npairs: number of total session-pairs included; Pooled CCC: Lin’s CCC pooled across (subject, session-pair, ROI, stimulus) combination; CI low: 95% CI lower limit; CI high: 95% CI upper limit.

| Condition | U | N | Npairs | Pooled CCC | CI low | CI high |
| --- | --- | --- | --- | --- | --- | --- |
| 2\|3\|25 | 1 | 2 | 1 | 0.672 | 0.672 | 0.672 |
| 2\|3\|50 | 3 | 7 | 5 | 0.623 | 0.381 | 0.715 |
| 2\|3\|100 | 1 | 2 | 1 | 0.670 | 0.670 | 0.670 |
| 4\|3\|50 | 2 | 4 | 2 | 0.710 | 0.661 | 0.760 |
| 8\|3\|25 | 2 | 5 | 4 | 0.590 | 0.525 | 0.676 |
| 8\|3\|50 | 7 | 26 | 40 | 0.608 | 0.478 | 0.674 |
| 8\|3\|100 | 2 | 5 | 4 | 0.549 | 0.515 | 0.611 |
| WC (8\|3\|50) | 5 | 17 | 40 | 0.588 | 0.429 | 0.662 |

The most robustly sampled classical retinotopy condition, 8|3|50 (7 subjects, 40 session-pairs), showed a pooled CCC of 0.61 (95% CI [0.478, 0.674]). The WC (8|3|50) condition, matched to the same underlying FWS parameters but drawn from the Word-Center paradigm stimuli, was comparably well sampled (5 subjects, 40 session-pairs) and showed a similar, slightly lower pooled CCC of 0.59 (95% CI [0.429, 0.662]).

Two conditions, 2|3|25 and 2|3|100, were each supported by only a single subject and a single session-pair, yielding CI intervals equal to the point estimate. Two additional FWS configurations (4|3|25 and 4|3|100), could not be evaluated at all, as no subject contributed more than one session under either configuration.

#### pRF Model Comparison: OG vs CSS

To assess how the choice of pRF model influences spatial estimates, we compared eccentricity values obtained with the OG and CSS models across all FWS configurations and ROIs.

OG-derived pRF center estimates were consistently more peripheral than those obtained with the CSS model across all FWS configurations and every targeted ROI in the three stimulus types (Fig. 13). The divergence was largest in higher-order areas (VO-1, V3a, IPS-0, IPS-1), reaching a 2.5° mean bias difference in several ROIs.

**Figure 13.**
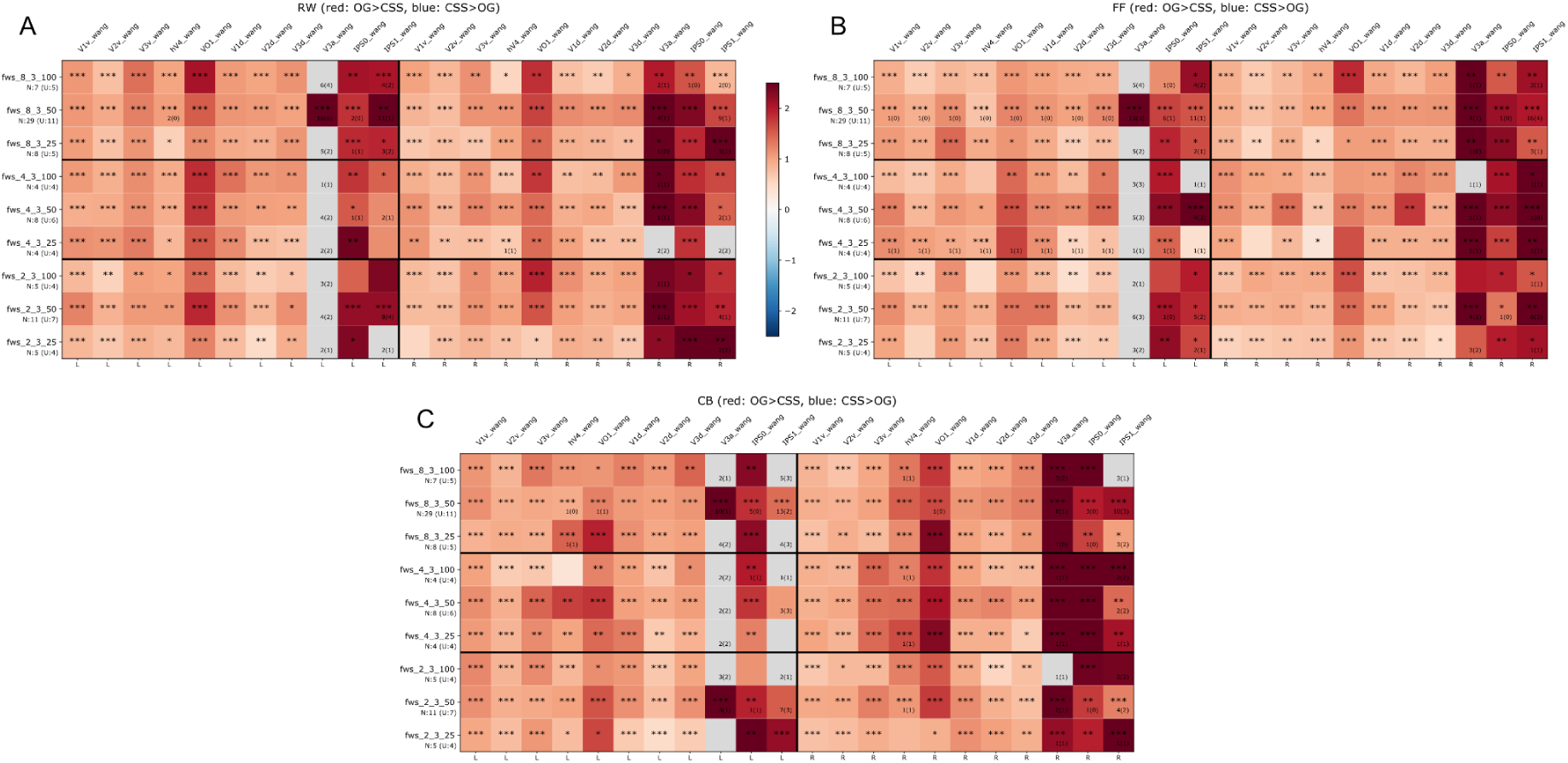
OG vs CSS eccentricity comparison matrices across all FWS configurations and visual ROIs. **A.** RW. **B.** FF. **C.** CB. Rows represent the nine FWS conditions. Columns represent the 22 ROIs from the Wang atlas; left-hemisphere ROIs occupy the first 11 columns and right-hemisphere ROIs the next 11, separated by a bold vertical line. Within each hemisphere, ventral-stream areas (V1v–VO1) precede dorsal-stream areas (V1d–IPS1), placing higher-order areas toward the right of each cluster. Cell color follows a diverging red/blue scale (±2.5°), showing mean eccentricity bias (OG − CSS, degrees of visual angle) between the paired model types: red indicates OG > CSS (more peripheral eccentricity for OG), blue indicates CSS > OG, and gray indicates no testable vertices. Significance was assessed per cell using a linear mixed-effects model of per-session-pair differences with a subject random intercept; stars mark cells where the null hypothesis of zero bias is rejected at p < 0.05 (***p < 0.001, **p < 0.01, *p < 0.05). Cells with fewer than two paired sessions or less than 25 paired-vertices were not tested. Row labels (N:x, U:y) indicate the number of contributing sessions (N) and unique subjects (U) for each FWS condition; per-cell deficit markers in the lower-right corner flag individual ROIs where the local sample falls below the row N(U) baseline.

### WC analyzed as retinotopy in visual and reading regions

#### WC carrier comparison

To characterize how the linguistic content and spatial arrangement of the fixated stimulus influence spatial tuning in visual ROIs, we compared pRF eccentricity estimates across the three WC conditions (fixRW, fixFF, and fixRWblock; Fig. 14).

**Figure 14.**
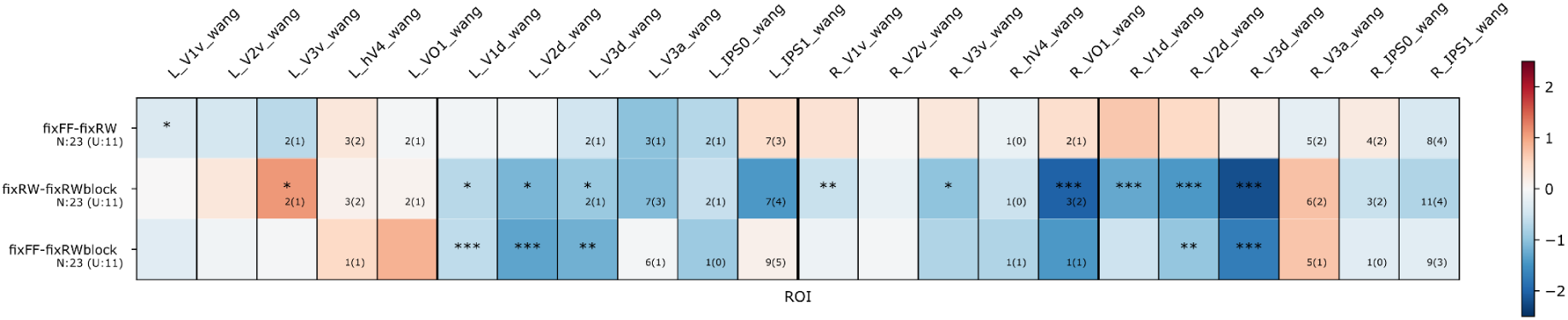
WC stimulus carrier comparison matrix across visual ROIs. Rows represent each WC stimulus pair (fixFF−fixRW, fixRW−fixRWblock, fixFF−fixRWblock). Columns represent the 22 ROIs from the Wang atlas; left-hemisphere ROIs occupy the first 11 columns and right-hemisphere ROIs the next 11, separated by a bold vertical line. Within each hemisphere, ventral-stream areas (V1v–VO1) precede dorsal-stream areas (V1d–IPS1), placing higher-order areas toward the right of each cluster. Cell color follows a diverging red/blue scale (±2.5°), showing mean eccentricity bias (Y − X, degrees of visual angle) between the paired stimuli types: red indicates Y > X (more peripheral eccentricity for the first stimulus), blue indicates X > Y, and gray indicates no testable vertices. Significance was assessed per cell using a linear mixed-effects model with a subject random intercept; stars mark cells where the null hypothesis of zero bias is rejected at p < 0.05 (***p < 0.001, **p < 0.01, *p < 0.05). Cells with fewer than two paired sessions or less than 25 paired-vertices were not tested. Row labels (N:x, U:y) indicate the number of contributing sessions (N) and unique subjects (U) for each FWS condition; per-cell deficit markers in the lower-right corner flag individual ROIs where the local sample falls below the row N(U) baseline.

A clear divergence in eccentricity estimates emerged between the continuous-string stimuli (fixRW and fixFF) and the blocked configuration (fixRWblock). In the left hemisphere, this took the form of a pathway-specific crossover: fixRW and fixFF produced similar or more peripheral responses than fixRWblock along the ventral pathway, whereas the dorsal pathway showed the opposite pattern, with fixRWblock yielding more peripheral eccentricity estimates than fixRW and fixFF.

In the right hemisphere, the stimulus-dependent effects were more uniform across the cortical hierarchy, with both ventral and dorsal pathways showing more peripheral responses for fixRWblock compared to fixRW and fixFF.

The comparison between fixRW and fixFF revealed no clear or systematic pattern of eccentricity differences across the visual and parietal hierarchies. Minor variations were observed, but these were characterized by inconsistent mean bias values, with one stimulus showing slightly higher eccentricity than the other across scattered ROIs without a discernible hierarchical or pathway-specific trend.

To determine whether the stimulus- and paradigm-dependent eccentricity effects observed in the visual cortex extend into the reading-selective cortex, we expanded the pRF eccentricity comparisons across the five vOTRC ROIs defined by the fLoc localizer (Fig. 15).

**Figure 15.**
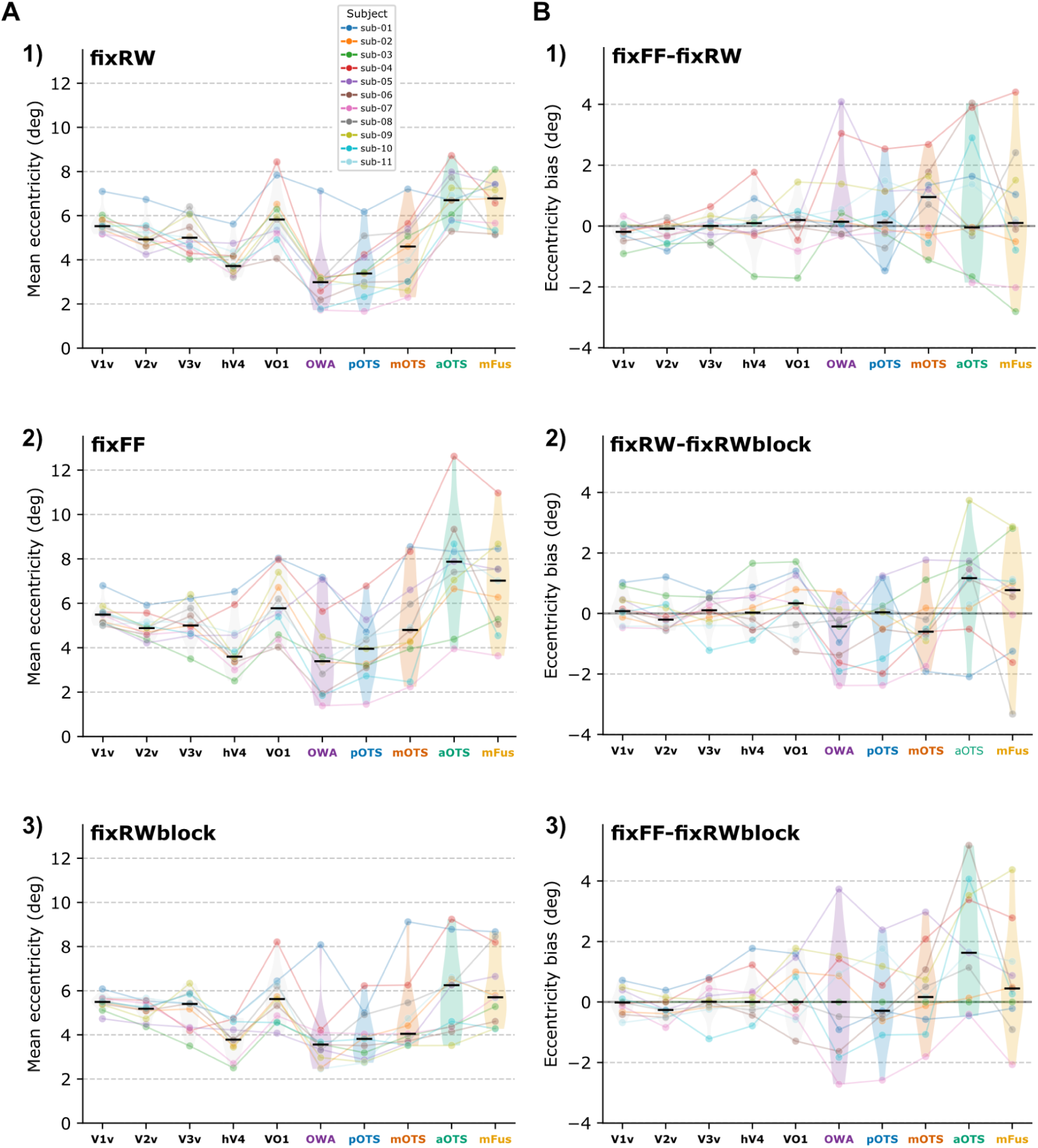
WC stimulus carrier comparison across all subjects and ventral visual and reading ROIs. Individual colored dots and connecting lines represent the 11 participants; black lines indicate the group median within each ROI. The X-axis tracks 10 ventral ROIs ordered from posterior to anterior. **A.** Y-axis shows the mean eccentricity per subject (in degrees). Rows display results for different carriers within the stimulus bar: 1) fixRW, 2) fixFF, and 3) fixRWblock. **B.** Y-axis shows the subject-specific eccentricity bias between carriers (in degrees). Rows display carrier differences: 1) fixFF−fixRW, 2) fixRW−fixRWblock, and 3) fixFF−RWblock.

For all three WC conditions considered individually (fixRW, fixFF, fixRWblock), mean eccentricity followed a similar shape across the gradient (Fig. 15, panel A). Across the early-to-mid ventral visual ROIs (V1v–V3v), eccentricity stayed relatively stable around 5°, dipped at hV4 (∼4°), and rose again at VO-1 (∼6°). Moving into the reading circuitry, eccentricity dropped sharply to its most foveal values anywhere in the gradient at OWA and pOTS (∼3–4°), before rising steeply through mOTS into aOTS and mFus, which showed the most peripheral estimates of any ROI in the gradient (∼6–8°).

Across-subject variability, visible as violin width, was consistently greater in the reading ROIs than in the more tightly clustered ventral visual ROIs.

The carrier-type mean bias plots showed a corresponding divide between the two regions (Fig. 15, panel B). Across the ventral visual pathway (V1v to VO-1), the median bias for all three contrasts remained close to zero, indicating no systematic carrier-type-dependent eccentricity shift in this portion of the gradient. Entering the reading circuitry, the fixFF−fixRW bias indicated a more foveal response to fixRW only in mOTS, while the rest of vOTRC ROIs showed no difference. Both fixRW-fixRWblock and fixFF−fixRWblock were close to zero or slightly negative in OWA, pOTS and mOTS, and positive in aOTS and mFus. As in panel A, across-subject variability in these bias estimates increased substantially moving into the reading circuitry, most visibly in aOTS and mFus.

#### Cross paradigm comparisons: classical retinotopy vs. Word-Center

To assess whether the concurrent presentation of sensory and linguistic information in the WC paradigm produces systematic shifts in pRF eccentricity relative to classical retinotopic mapping in visual ROIs, we compared eccentricity estimates between the two paradigms for stimulus pairs sharing at least one common component.

Contrasting CB stimuli against the three WC conditions (fixRW, fixFF, and fixRWblock) revealed a divergence in spatial tuning that was consistent only within the left hemisphere (Fig. 16). Along the ventral pathway, CB elicited increasingly higher pRF eccentricity estimates compared to the WC stimuli, with the most pronounced difference observed in area VO-1. Conversely, the dorsal pathway exhibited the opposite effect, with WC conditions producing increasingly higher eccentricity estimates than CB (largest difference observed in IPS-0 and IPS-1).

**Figure 16.**
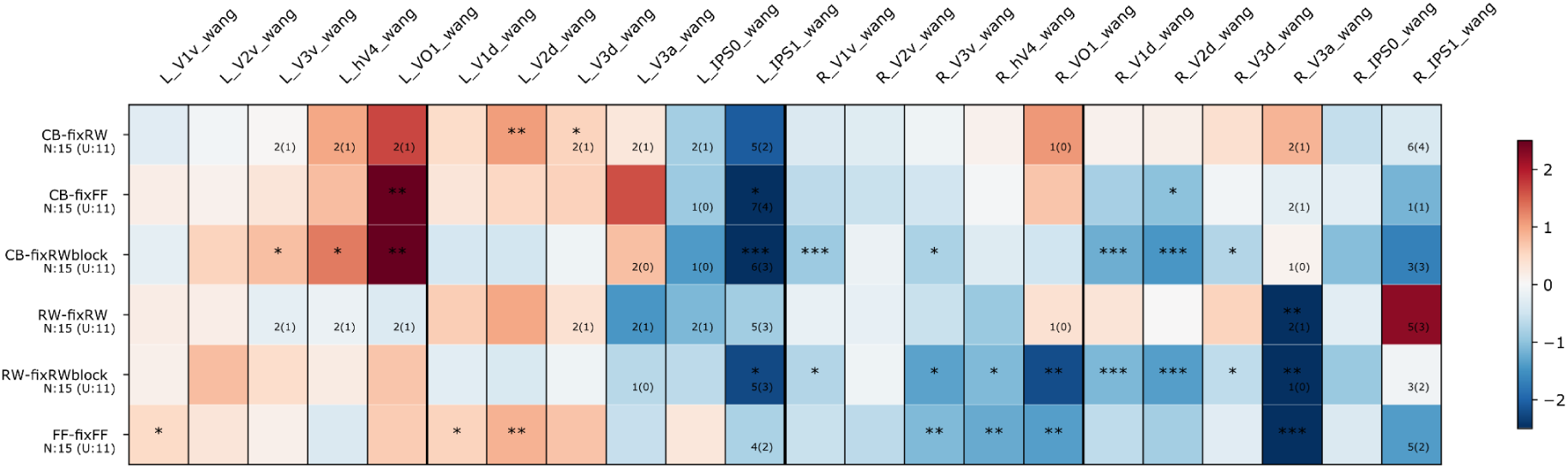
Classical retinotopy vs WC stimulus carrier comparison matrix across visual ROIs. Rows represent each classical retinotopy−WC stimulus pair (CB−fixRW, CB−fixFF, CB−fixRWblock, RW−fixRW, RW−fixRWblock, FF−fixFF). Columns represent the 22 ROIs from the Wang atlas; left-hemisphere ROIs occupy the first 11 columns and right-hemisphere ROIs the next 11, separated by a bold vertical line. Within each hemisphere, ventral-stream areas (V1v–VO1) precede dorsal-stream areas (V1d–IPS1), placing higher-order areas toward the right of each cluster. Cell color follows a diverging red/blue scale (±2.5°), showing mean eccentricity bias (Y − X, degrees of visual angle) between the paired stimuli types: red indicates Y > X (more peripheral eccentricity for the first stimulus), blue indicates X > Y, and gray indicates no testable vertices. Significance was assessed per cell using a linear mixed-effects model with a subject random intercept; stars mark cells where the null hypothesis of zero bias is rejected at p < 0.05 (***p < 0.001, **p < 0.01, *p < 0.05). Cells with fewer than two paired sessions or less than 25 paired-vertices were not tested. Row labels (N:x, U:y) indicate the number of contributing sessions (N) and unique subjects (U) for each FWS condition; per-cell deficit markers in the lower-right corner flag individual ROIs where the local sample falls below the row N(U) baseline.

In the right hemisphere, these pathway-specific opposing effects were absent: WC stimuli yielded higher eccentricity estimates than classical retinotopic stimuli across most ROIs, in both the ventral and dorsal pathways.

A similar, though weaker, ventral-dorsal divergence was observed for the RW–fixRWblock comparison in the left hemisphere, with RW yielding more peripheral responses along the ventral pathway and fixRWblock yielding more peripheral responses along the dorsal pathway, mirroring the pattern seen for CB but at a reduced magnitude.

To assess whether the eccentricity shifts observed with classical retinotopy remain when using the Word-Center (WC) paradigm, we conducted the same cross-paradigm comparisons across the 10 left ventral ROIs (5 visual: V1v to VO-1; 5 reading: OWA to mFus) ordered along a posterior-to-anterior gradient (Fig. 17). Because the WC stimuli utilized an 8|3|50 FWS configuration, all comparisons are paired with classical retinotopy sessions acquired with identical parameters, resulting in paired intersession measurements. CB vs. fixRW and fixFF showed similar eccentricities across early visual ROIs, and in the reading regions differences emerged with no clear pattern (Fig. 17, panels A and E).

**Figure 17.**
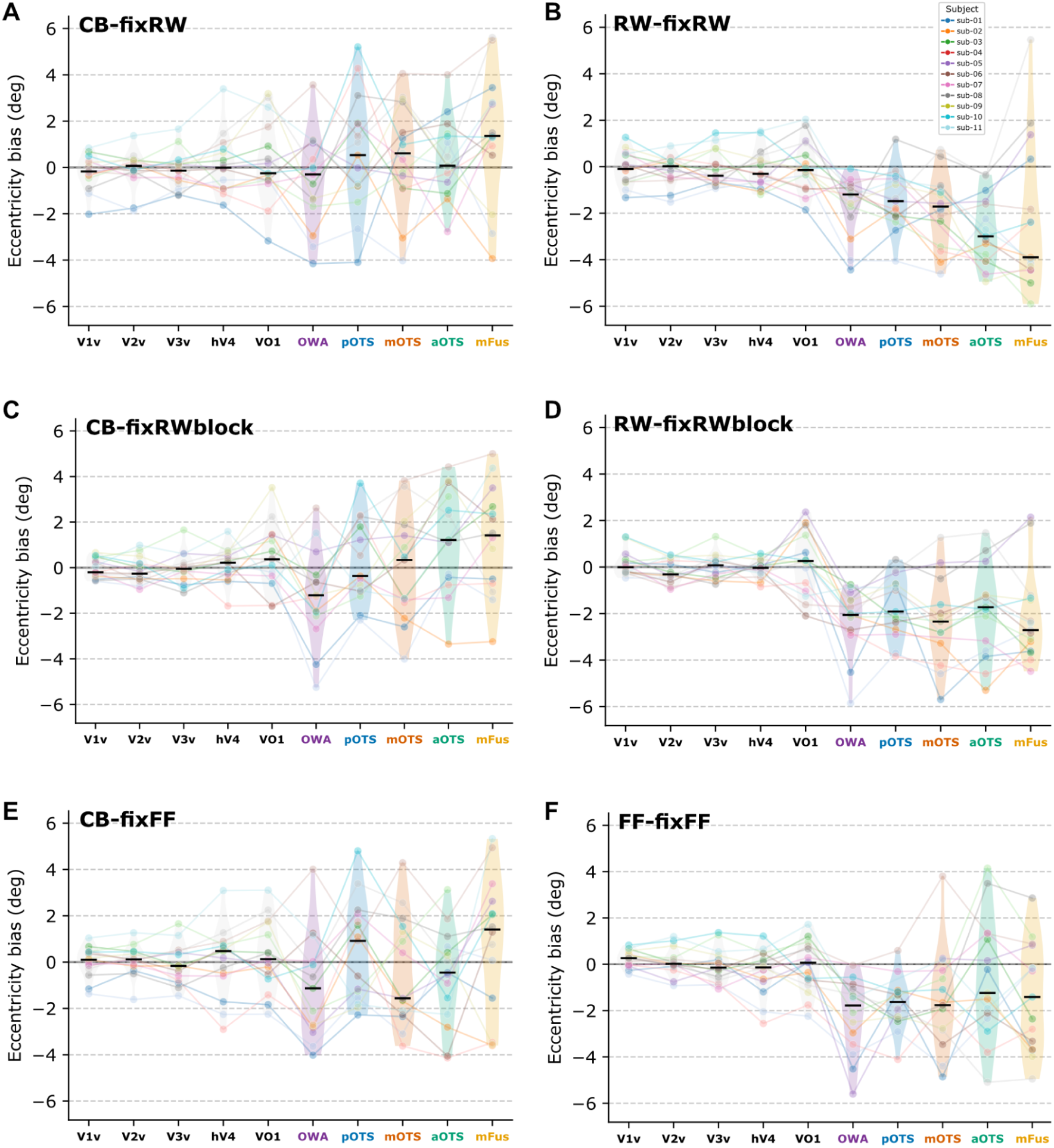
Classical retinotopy vs WC stimulus carrier comparison across all subjects and ventral visual and reading ROIs. Individual colored dots and connecting lines represent the 11 participants; horizontal black ticks indicate the group median within each ROI. The X-axis displays 10 ventral ROIs (5 visual: V1v–VO1; 5 reading: OWA–mFus) ordered along a posterior-to-anterior gradient. Y-axis shows the subject-specific eccentricity bias between carriers in degrees of visual angle. Individual panel titles denote the specific carriers compared. **A.** CB−fixRW. **B.** RW−fixRW. **C.** CB−fixRWblock. **D.** RW−fixRWblock. **E.** CB−fixFF. **F.** FF−fixFF.

On the other hand, the CB vs. fixRWblock comparison reveals a clear divergence in spatial tuning between pathways: along the ventral visual path and the reading regions, CB yields increasingly more peripheral estimates than the WC stimulus. In both cases, the most posterior ROIs (V1v and V2v, OWA and pOTS) start with slightly more foveal estimates for CB, while the most anterior ones show slightly more foveal (hV4 and VO-1) and considerably more foveal (aOTS and mFus) responses for fixRWblock (Fig. 17, panel C).

The other three comparisons, RW vs. fixRW, RW vs. fixRWblock, and FF vs. fixFF (Fig. 17, panels B, D and F), exhibit a consistent pattern: minimal differences in visual regions but systematically higher eccentricity values for WC stimuli across the reading hierarchy.

### WC analyzed as word localizer in reading regions

To assess whether the fixRWblock condition reliably engaged the reading-selective cortex, we examined the RWvsNull contrast T-values within the five fLoc-defined reading ROIs. Group median T-values differed substantially across regions, following a posterior-to-anterior gradient (Fig. 18): the highest mean responses were observed in the more posterior OWA (median T = 11.7) and pOTS (median T = 11.6) ROIs, along with mOTS (median T = 10.9), while the most anterior regions (aOTS: median T = 4.7; mFus: median T = 5) showing low group-mean responses.

**Figure 18.**
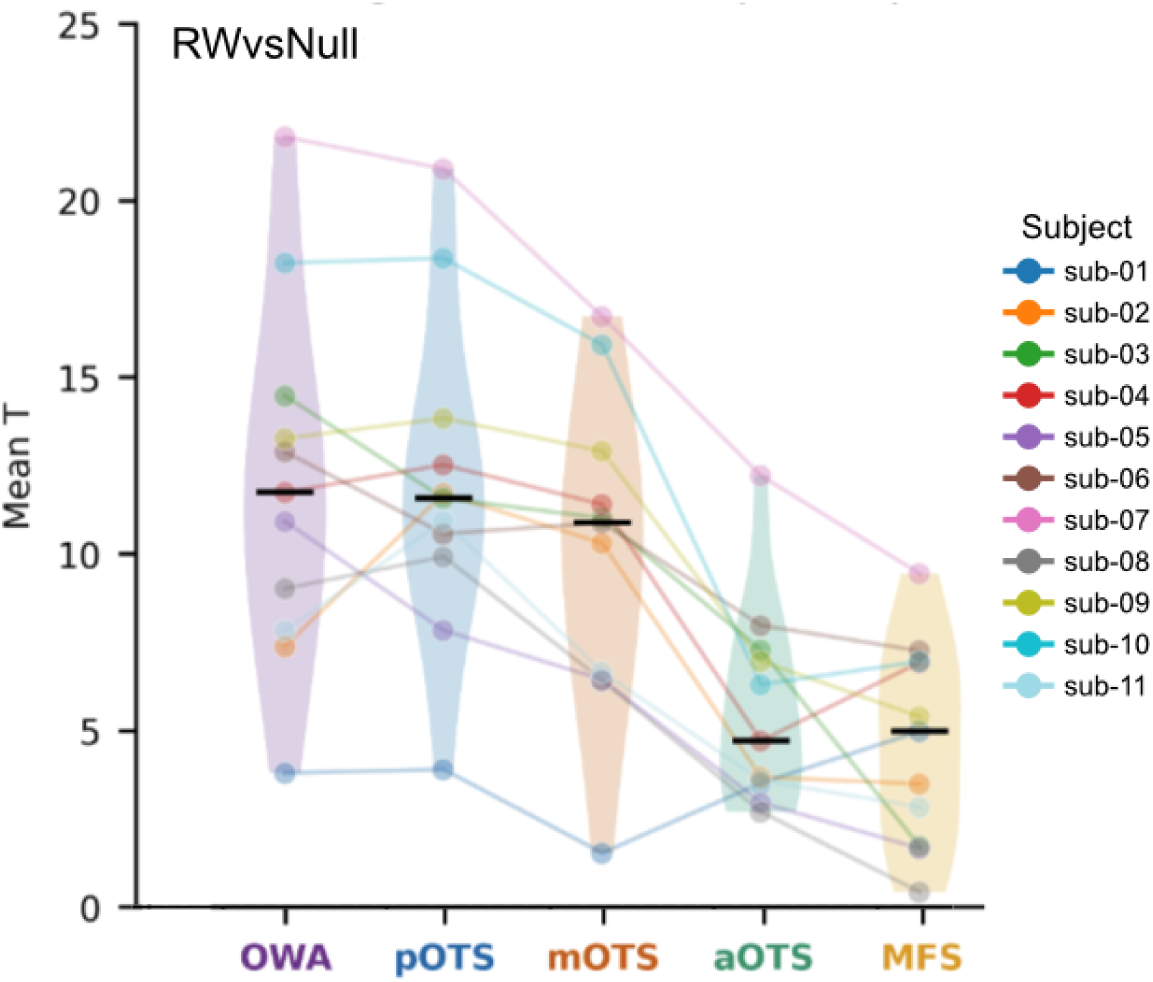
Word-selectivity localizer contrast (RWvsNull) T-value distribution across reading-selective ROIs. Group- and subject-level T-values for the RWvsNull contrast, used to assess how reliably the fixRWblock condition engaged reading-selective cortex, are shown for the five fLoc-defined vOTRC ROIs, arranged along the x-axis from the most posterior (OWA) to the most anterior (mFus, labeled MFS) region. Violin shapes show the across-subject distribution of mean T-values within each ROI, with the horizontal black tick marking the group median. Each colored, semi-transparent line connects a single subject’s (n = 11) mean T-value across the five ROIs, with matching colored dots marking that subject’s value in each region; subject-to-color correspondence is given in the legend (sub-01 to sub-11).

Considerable across-subject variability was present in all five ROIs. This was most pronounced in OWA, pOTS and mOTS, where individual subject T-values ranged broadly despite comparably high group means. aOTS and mFus showed lower group means overall, with several subjects in these regions showing near-null or minimal T-values, alongside a small number of individuals with comparatively high responses.

## Discussion

The primary aim of this study was to establish a rigorous sensory baseline within the visual reading hierarchy, a necessary prerequisite for isolating the top-down cognitive signals described by the Interactive Account (Price & Devlin, 2011). By systematically evaluating stimulus-dependent pRF shifts and validating the novel Word-Center (WC) paradigm, this study addressed the intense methodological challenges inherent in mapping the transition from low-level vision to abstract linguistic processing.

Our findings provide a robust individual-level replication of hierarchical spatial tuning shifts, where orthographic carriers elicit systematically more foveal estimates than checkerboards, while highlighting the critical role of FWS (Frequency, Width, and Size) parameters in achieving parameter stability. Furthermore, the validation of the WC paradigm demonstrates its efficacy in decoupling bottom-up sensory sweeps from central cognitive processing, serving as a dual-purpose tool for simultaneous retinotopy and functional localization. Together, these results offer a foundational framework for a more precise, multi-signal characterization of the human reading circuitry.

### Classical retinotopy

#### Stimulus carrier comparisons

Our FWS- and subject-level results show that pRF eccentricity differences between orthographic stimuli (RW, FF) and checkerboards increase progressively along the visual hierarchy, with the largest divergence in higher-order areas of the ventral stream. In this pathway, the pattern (RW producing systematically more foveal pRF estimates than CB, growing more pronounced toward anterior maps) directly replicates the hierarchical stimulus-dependency reported by Lerma-Usabiaga et al. (2021), and fits well with the broader literature on the functional organization of high-order visual cortex. Hasson et al. (2002) showed that letter strings and words, unlike object categories such as buildings, are preferentially represented within the foveally-biased portion of high-order ventral cortex, consistent with the idea that regions specialized for tasks demanding fine visual discrimination, such as reading, are organized around a central visual field bias (Hasson et al., 2002; Nordt et al., 2019). Under this account, RW stimuli would be expected to more strongly recruit the foveally-tuned subpopulation of vertices within higher-order ventral regions than a low-level control stimulus like CB, producing precisely the kind of anterior-weighted, foveally-shifted bias we observe. As this analysis is restricted to purely sensory regions prior to the vOTRC, this difference in eccentricity does not appear in the perceptually similar but cognitively distinct stimuli (RW and FF), where the eccentricity bias is small along the ventral pathway.

Crucially, the results in panel B of Fig. 10 provide a replication of the reading effect previously reported in Lerma-Usabiaga et al. (2021). While FF and RW produce nearly identical eccentricity estimates in early visual areas and posterior reading ROIs (OWA, pOTS), a significant divergence occurs in the aOTS. In this region, FF and CB converge with practically no difference in eccentricity, whereas RW stimuli elicit a highly distinct foveal shift. This suggests that while posterior regions treat orthographic strings primarily as visual features, consistent with the role attributed to pOTS in parallel feature extraction (White et al., 2019; Caffarra et al., 2021), the aOTS isolates true lexical items, treating visually matched false-fonts as equivalent to low-level checkerboard patterns.

In the dorsal pathway, the picture is less clear. Higher-order dorsal areas (V3a, IPS-0, IPS-1) do show larger-magnitude eccentricity biases than earlier dorsal areas (V1d, V2d, V3d), but, unlike the ventral pathway, this bias is not consistently showing CB yielding more peripheral estimates than RW and FF in these regions. Much of this pattern is also difficult to interpret at face value, since a large proportion of cells in these higher-order dorsal ROIs, particularly V3a, lack sufficient vertex pairs to be tested at all, a consequence of the sparse and variable coverage these regions receive under the Wang et al. (2015) atlas (see Limitations). Given this, we cannot currently distinguish whether the larger biases in higher-order dorsal cortex reflect a genuine form of stimulus sensitivity in this pathway, or whether they are largely an artifact of the sparse and uneven sampling in these regions. Denser or individually defined dorsal ROIs would be needed to properly judge between these possibilities before any significant interpretation can be drawn for the dorsal stream.

At the subject level, the hierarchical ventral pattern was present in 70% of participants, a proportion broadly consistent with the approximately 80% reported by Lerma-Usabiaga et al. (2021). The reason why some subjects did not show the pattern at all is unclear, but one possible explanation follows: because reading is a learned skill, the cortical territory that comes to support it is thought to be individually shaped, or “recycled”, from more general-purpose visual cortex through each person’s own reading experience (Dehaene & Cohen, 2007), a process already linked to substantial individual variability in the location and tuning of nearby word-selective cortex (Glezer & Riesenhuber, 2013). If hV4 and VO-1, the regions immediately preceding the reading circuitry, are shaped by this same individually variable process, a group-based ROI atlas (Wang et al., 2015) may not always align precisely with a given participant’s true functional boundaries, potentially obscuring the effect in some subjects. A natural next step for our data would be to characterize these ROIs individually per subject and session, to test whether this improves the consistency of the effect across participants.

Beyond individual anatomical variability, the 30% of subjects who did not show the expected hierarchical shift may be influenced by the stability of specific FWS configurations. As our results indicate that pRF estimates are sensitive to flickering frequency and element size, certain parameter combinations may inadvertently attenuate these spatial effects in specific individuals, highlighting the importance of standardized, high-stability FWS parameters for individual-subject applications.

#### Stimulus frequency and size effect

The frequency manipulation produced a clear and systematic bias, with lower temporal frequencies (2 Hz) yielding more peripheral eccentricity estimates than higher frequencies (8 Hz) at element sizes 25 and 50. This effect disappeared at the largest element size (100), pointing to an interaction between the stimulus’s temporal and spatial properties rather than an independent effect of frequency alone. The size manipulation, by contrast, produced a weaker and less uniform bias, present mainly for the largest size contrasts (25 vs. 50 and 25 vs. 100) at lower frequencies, and largely attenuated at 8 Hz.

Taken together, both effects appear only under specific combinations of the other parameter, rather than holding uniformly across the full FWS space: the frequency effect vanished at the largest element size, and the size effect was only clearly present at lower frequencies. This suggests that temporal frequency and element size do not act independently, but jointly determine the effective stimulus drive at a given retinal location, meaning a given flicker rate may only bias pRF estimates within a certain range of element sizes, and vice versa.

Beyond the specific mechanism, these results reinforce a theme recurring throughout this study: pRF estimates are not a neutral readout of neural spatial tuning but are shaped by choices made well upstream of model fitting, including the low-level temporal and spatial properties of the mapping stimulus itself. This has practical implications for pRF studies of category-selective cortex, where comparing pRF estimates across stimulus conditions requires these low-level properties to be matched in order to avoid attributing what may be an incidental low-level bias to a genuine category-selective effect. Given the exploratory nature and limited session counts underlying this particular analysis, we treat it as a motivation for future, purpose-built studies rather than a settled account of how temporal and spatial stimulus properties shape pRF estimates.

#### pRF eccentricity reliability

Test-retest reliability, assessed via Lin’s CCC on paired vertex-level eccentricity estimates, was consistently positive but moderate across the eight tested FWS configurations, with pooled CCC values ranging from 0.55 (8|3|100) to 0.71 (4|3|50) (Table 1). None of the conditions approached the high end of the CCC scale, indicating that session-to-session agreement in pRF eccentricity, while systematic, is far from perfect. The two most robustly sampled conditions, classical retinotopy 8|3|50 (7 subjects, 40 session-pairs) and the matched WC 8|3|50 condition (5 subjects, 40 session-pairs), yielded similar pooled CCCs of 0.61 and 0.59 respectively, suggesting that the WC paradigm does not introduce any additional instability in pRF eccentricity estimation relative to classical retinotopic mapping, and that the moderate reliability observed here is likely a general property of our pRF fitting pipeline rather than a paradigm-specific issue.

This level of reliability is directly relevant to how the stimulus- and paradigm-dependent effects should be interpreted. A moderate CCC indicates that a non-trivial share of the variance in vertex-level eccentricity estimates reflects session-to-session measurement noise rather than genuine, stable differences in pRF position. Because our mean-bias comparisons rely on across-session pairings, this noise should work against, rather than in favor of, detecting the stimulus-dependent shifts we report; the fact that several comparisons nonetheless reached significance despite this added noise floor increases our confidence that the underlying effects are genuine, even if their exact magnitude is likely attenuated by measurement error.

Given how unbalanced and multi-layered our data structure is, pooling across subjects, session-pairs, ROIs and stimulus types was considered the best way to obtain a stable summary number for interpretation. Nevertheless, this comes at a cost: each median discards information about how reliability varies across subjects, ROIs, and stimuli, collapsing what is likely a genuinely heterogeneous pattern, for instance the ROI-level instability for V3a, IPS-0, and IPS-1 (see Limitations), into a single point estimate per FWS configuration. The pooled CCCs in Table 1 should therefore be read as a coarse, conservative summary of reliability rather than a precise estimate of it.

As this loss of information comes from the unbalanced structure of our data, rather than from any shortcoming of the CCC metric itself, we do not think a different classical reliability statistic would resolve it. We are therefore exploring a hierarchical Bayesian modeling (HBM) approach as a more principled alternative for future work. Rather than aggregating cell-level estimates through a fixed cascade of medians, an HBM would model sessions as nested within subjects, subjects within FWS configurations, and estimates within ROIs directly, allowing sparser cells to borrow statistical strength from better-sampled ones through partial pooling instead of being excluded or collapsed outright. This would let us retain, and explicitly quantify, variability at each level of the data structure, including between-subject and between-ROI differences in reliability that the current pooled-CCC approach cannot recover, while still yielding principled uncertainty estimates for the sparser FWS configurations that currently produce single-point or entirely untestable CCCs.

#### pRF model comparison: OG vs CSS

The finding that OG modeling systematically yields more peripheral eccentricity estimates than CSS across all FWS configurations and ROIs is consistent with the known consequences of unmodeled subadditive spatial summation in human visual cortex. Kay et al. (2013) showed that linear models like OG can overestimate pRF size under compressive, subadditive responses: because the OG model assumes a purely linear relationship between stimulus extent and response amplitude, it cannot account for the fact that a vertex’s response to a large stimulus is often smaller than the sum of its responses to the stimulus components, and must instead recruit a larger Gaussian to approximate this saturating pattern. While Kay et al. (2013) reported this effect specifically for pRF size, we propose that this same mechanism plausibly extends to pRF center position: a compensatory, oversized Gaussian may also shift its fitted center away from the true receptive field location, offering a possible explanation for the more peripheral OG estimates observed in our results.

This account is supported by the spatial pattern of the effect. The OG-CSS divergence was smallest in early visual areas and grew progressively larger in higher-order regions of both ventral and dorsal pathways (VO-1, V3a, IPS-0, IPS-1), consistent with reports that subadditive summation intensifies along the visual hierarchy as pRFs grow larger and more overlapping (Kay et al., 2013; Le et al., 2017; Lerma-Usabiaga et al., 2021). These are precisely the regions where a linear model has the most unmodeled nonlinearity to compensate for, and thus the greatest opportunity for its spatial estimates to be distorted.

These results suggest that model choice is not merely a technical preference but a factor that can meaningfully influence the absolute spatial boundaries assigned to the visual cortex. Following the proposed link between subadditive summation and eccentricity bias, a nonlinear framework like CSS would offer a more biologically plausible account of spatial tuning in these areas. Confirming this link and exploring how different models affect pRF parameters is an important direction for future work. From the perspective of the Interactive Account, the choice of a biologically plausible model like CSS is not merely a technical preference but a prerequisite. Because linear models (OG) systematically yield more peripheral estimates in higher-order regions, failing to account for these nonlinearities could lead to the misattribution of a purely sensory model-distortion as a top-down cognitive modulation.

### Word-Center paradigm

#### WC stimulus carrier comparison in visual and reading regions

The lack of a systematic eccentricity difference between fixRW and fixFF across all analyzed ROIs indicates that the visual hierarchy, prior to entering the vOTRC, does not differentiate between centrally presented RW and visually matched FF. This aligns with the sensory model of early and extrastriate visual maps, where responses are driven by low-level visual features rather than the symbolic or lexical identity of the stimulus (Lerma-Usabiaga et al., 2021). Because these regions do not yet integrate the cognitive signals necessary to separate lexical items from visually similar non-lexical strings, they treat both orthographic types as equivalent sensory inputs.

A different pattern emerges when comparing the orthographic strings (fixRW/fixFF) to the blocked condition (fixRWblock). In the right hemisphere, both the ventral and dorsal pathways showed more peripheral pRF estimates for fixRWblock than for the continuous strings. Because reading is a strongly left-lateralized skill (Cohen et al., 2000; Gaillard et al., 2006), the right hemisphere is thought to play a comparatively minor role in specialized linguistic processing of centrally presented orthographic content (Vinckier et al., 2007; Glezer and Riesenhuber, 2013). Under this view, what matters is simply how much of the time a foveal stimulus is present at all: in fixRW and fixFF, a word or false-font string occupies fixation continuously, whereas in fixRWblock it is present only intermittently, leaving a larger share of the run without any foveal stimuli. Therefore, we suggest that, with less consistent stimulation competing against the peripheral sweep of the bars, the fitted pRF centers in fixRWblock are pulled comparatively further into the periphery.

In the left hemisphere, the dorsal pathway showed this same pattern, which fits well with the established role of the IPS in representing spatial attention and priority, a function that is not itself specialized for reading and operates similarly across both hemispheres (Yeatman & White, 2021). On the other hand, the left ventral stream was the only pathway to depart from this pattern, showing no difference or even more eccentric estimates for the continuous strings compared to fixRWblock. Since this is the only pathway-hemisphere combination that eventually terminates in the reading-specific vOTRC, we propose that a second, competing influence is at play here: alongside the same general foveal pull described above, the ventral stream’s role in processing the spatial horizontally elongated structure of orthographic strings may introduce an opposing peripheral pull that scales with how often such strings are presented. Under this account, fixRW and fixFF, presented continuously, are subject to the strongest peripheral pull, while fixRWblock, with strings appearing only intermittently, is subject to comparatively little of it, resulting in more foveal estimates. This two-factor account, balancing a general foveal pull with a stimulus-specific peripheral shift, is further supported by the habituation effect identified in our methodology (see WC localizer values in reading regions). The constant presentation of orthographic material in the fixRW condition likely induced neural fatigue, resulting in the higher noise levels and the more eccentric pRF estimates observed in the left ventral stream compared to the intermittent fixRWblock design.

While these ventral ROIs are not part of the reading circuitry itself, the pathway-specific differences in how they encode strings versus blocks may be constrained by pre-existing structural connectivity via the AF that prepares the left hemisphere for the high-acuity, parallel processing required for reading (Lerma-Usabiaga et al., 2018; White et al., 2019; Yablonski et al., 2024). These results suggest that while the “reading” distinction is not yet present, the computational groundwork for processing the structural properties of text is already being differentiated at these earlier stages of the visual hierarchy (Lerma-Usabiaga et al., 2021).

Extending this comparison to the reading-selective vOTRC (Fig. 15) revealed a pattern that fits well with a lexical, rather than purely sensory, account. Unlike the visual ROIs, where fixRW and fixFF produced largely indistinguishable pRF estimates, only the mOTS region showed a more foveal response to fixRW than to fixFF. Because the two stimulus types are matched for low-level visual properties and differ only in their lexical status, we interpret this foveal bias for real words as reflecting the additional word-recognition processing that this region, unlike all others, performs on the fixated string. When comparing, fixFF and fixRWblock, while, once again, the visual regions produced largely indistinguishable pRF estimates, only aOTS (and to a lesser extent, mFus) showed a more foveal response to fixRW than to fixFF, even when the stimulus with false-fonts stays in the center during a higher amount of time. These results are consistent with prior evidence that anterior regions in the OTS, rather than upstream visual cortex, are the stage at which real words are distinguished from visually matched non-lexical strings such as false-fonts (Lerma-Usabiaga et al., 2021; Vinckier et al., 2007).

This perceptual-lexical divergence reinforces the functional segregation of the vOTRC described above (see Functional Segregation and Sensory-Cognitive Integration in the vOTRC), identifying pOTS as the boundary where sensory feature extraction dominates over lexical modulation (Lerma-Usabiaga et al., 2018; White et al., 2019; Caffarra et al., 2021). This could bring an explanation as to why mOTS and aOTS showed more foveal responses to RW stimuli compared to FF, while pOTS showed no difference. However, it is still unclear why the expected bias was restricted to the fixFF and fixRW comparison in mOTS, while in aOTS it only appeared when comparing fixFF to fixRWblock.

#### Cross-paradigm comparisons: classical retinotopy vs. Word-Center in visual and reading regions

Contrasting CB against the three WC conditions supports the two-factor account proposed above. In the left ventral pathway, CB elicited the most peripheral pRF estimates when compared to the three WC stimulus types. This fits the proposed account directly: as the central stimuli is not present in CB, there is no foveal pull and pRF centers are much more peripheral than those from the three WC stimuli types.

In the left dorsal pathway, the pattern reverses: CB elicited the most foveal estimates of the four conditions in IPS-0 and IPS-1, while all three WC conditions produced more peripheral estimates. The mechanism underlying this result is not yet clear. Unlike the within-WC comparisons above, where the relevant manipulation was how often central content was presented, the key difference here is simply between having a linguistic task at fixation at all (any WC condition) versus not (CB), and this cannot be explained by a difference in fixation demand, since both CB and the WC conditions required participants to maintain central fixation throughout. This pattern is also present across most ROIs of the right hemisphere. One tentative possibility is that actively attending to and reading linguistic content at fixation, even intermittently, recruits attentional or cognitive engagement beyond what simple fixation requires, and that this additional engagement is reflected in the spatial profile of the IPS response in a way that the pRF model captures as a more peripheral estimate. This remains an open question that we aim to address in future work, using eye-tracking or explicit manipulations of cognitive load at fixation to isolate what aspect of the WC conditions drives this shift. Furthermore, the observed spatial shifts in the IPS may be constrained by underlying structural connectivity, specifically the Vertical Occipital Fasciculus (VOF), which facilitates direct communication between the dorsal attention network and the pOTS (Yablonski et al., 2024; Weiner et al., 2017b). This suggests that the peripheral shifts observed in the IPS during linguistic tasks may reflect the recruitment of this specific structural circuit

Extending this comparison into the reading circuitry (Fig. 17) revealed a more differentiated picture. CB vs. fixRW and CB vs. fixFF showed no consistent pattern in either the visual or reading portions of the gradient. The CB vs. fixRWblock comparison, however, was the one case where visual and reading regions showed a genuinely convergent effect: CB produced increasingly more peripheral estimates than fixRWblock moving anteriorly through both the left ventral visual pathway and the reading circuitry. Because CB and fixRWblock share the same checkerboard carrier and differ only in whether words also intermittently occupy fixation, this comparison should largely cancel out the shared low-level sensory drive common to both stimuli, isolating the contribution of the central linguistic content itself. Read this way, the systematically more foveal response to fixRWblock suggests that, of the three WC conditions, fixRWblock is the one that most reliably engages top-down linguistic processing strongly enough to pull pRF estimates toward fixation, over and above whatever the checkerboard carrier contributes on its own.

A different, and considerably harder to interpret, pattern emerged for the remaining three comparisons, RW vs. fixRW, RW vs. fixRWblock, and FF vs. fixFF. In all three, the visual regions showed minimal difference between paradigms, but WC stimuli produced systematically more peripheral estimates than their classical retinotopic counterparts throughout the reading circuitry. We do not have a confident explanation for this pattern, but one candidate difference between the paradigms may be relevant: the classical retinotopic bar stimuli present a dense string of multiple RW or FF items filling the width of the bar, whereas the corresponding WC stimuli present a single word or false-font in isolation at fixation. As the multi-item string inside the bars sweeps through central vision, if it more closely approximates the visual structure of continuous reading than a single isolated item does, this could plausibly elicit a more foveal response than the single fixated WC stimulus produces at fixation. We offer this as a tentative hypothesis rather than a confirmed account, and see it as a natural target for follow-up work, for instance by parametrically varying the number of simultaneously presented items in a fixated stimulus.

#### WC localizer values in reading regions

The clearest pattern in these results is a posterior-to-anterior gradient in how robustly fixRWblock drove the fLoc-defined reading ROIs: posterior regions (OWA, pOTS, and mOTS) showed strong group-mean responses, while the more anterior regions (aOTS and mFus) showed considerably weaker mean responses. This is consistently observed in our previous work with localizers and in the fLoc data analysis as well, where all subjects show a posterior to anterior gradient in T-values (Lerma-Usabiaga et al., 2026).

These results support a multi-patch/gradient account of the reading network (Lerma-Usabiaga et al., 2018; Yablonski et al., 2024) over a classic, single-module VWFA (Cohen et al., 2000; Dehaene & Cohen, 2011). By demonstrating that adjacent patches such as OWA, pOTS, and mOTS exhibit distinct functional profiles and varying signal-to-noise ratios, this analysis validates the necessity of the high-resolution, multi-ROI approach adopted throughout this study.

Finally, this analysis was intended as a first-pass validation rather than a rigorous statistical test of localizer suitability, and it has some clear limitations as such. We did not apply a formal per-subject or per-ROI threshold (e.g., a minimum T-value, as is common practice when validating functional localizers) to quantify what proportion of subjects meaningfully “pass” as showing a reading-selective WC response in each ROI; doing so, particularly at a conventional threshold, would likely reveal that a non-trivial fraction of subjects fall below it in aOTS and mFus specifically. Future work should apply such a threshold explicitly, and would also benefit from individually defined reading ROIs, rather than the group fLoc atlas used here, to determine whether the weaker and more variable responses in anterior regions reflect a genuine limitation of the fixRWblock stimulus or, at least in part, an artifact of imprecise group-level ROI localization.

### Limitations

A primary limitation of the present study is its predominant focus on retinotopic analyses. The reading-selective ROIs identified by the fLoc localizer were not directly characterized using the WC paradigm results at the individual-subject level. Future work should implement an automated ROI definition approach (autoROI) to isolate these functional clusters in each participant individually, moving beyond predefined anatomical or group-based benchmarks, and use Dice coefficient analyses to quantitatively evaluate the reliability and spatial specificity of these newly localized regions against the established localizer results. Such an approach would help determine whether the WC paradigm can serve as a sensitive, independent tool for dissociating the sensory and cognitive contributions to word recognition.

Although the dense-sampling design (11 participants × 10 sessions) maximizes within-subject signal-to-noise ratio and mapping stability, the small number of participants limits the generalizability of these findings to broader populations, including left-handed individuals or those with varying levels of reading proficiency. The sample was also linguistically heterogeneous, comprising native speakers of five different languages (Spanish, Austrian German, Italian, French, and Mandarin Chinese), with stimuli adapted accordingly for each language, which may introduce additional variability in reading-related responses that was not explicitly modeled. In sum, the range of FWS configurations used across sessions was not uniform within or across participants, which introduces an additional source of variability in cross-session comparisons and reduces the number of available session pairs sharing identical configurations. This heterogeneity has direct analytical consequences: in the Subject-level stimulus carrier comparison analysis, sessions were pooled across FWS configurations to maximize within-subject pairings, meaning that the resulting variability reflects not only carrier-type effects but also differences in temporal frequency and element size between paired sessions; at the FWS level, conversely, restricting comparisons to session pairs sharing identical configurations reduced the number of available pairs, constraining the statistical robustness of those analyses. Future work should replicate this paradigm with a larger and more demographically diverse sample while standardizing FWS configurations across sessions to increase comparison power.

Despite using the FWS (flickering frequency, bar width and element size) acronym, we did not manipulate the width of the bar for this experiment (as the three 4|4|26 sessions were excluded for analysis). This variable was excluded from the design because manipulating it in addition to frequency and element size would have produced an unmanageably large number of stimulus combinations given the available scanning time. Nevertheless, bar width may substantially affect the results, and we recommend investigating this parameter in future work.

A further limitation concerns the reliability of eccentricity estimates in several of the dorsal visual ROIs, particularly V3a, IPS-0, and IPS-1. Because ROI boundaries were defined using the Wang et al. (2015) probabilistic atlas, which is derived from group-level retinotopic maps, areas with high inter-individual anatomical variability tend to yield fewer vertices when the atlas is projected onto individual anatomy. This results in a substantially lower number of vertex pairs available for cross-session comparisons in these regions, making the corresponding estimates less stable and harder to interpret with confidence, with V3a being the most affected. To mitigate the risk of drawing conclusions from insufficiently sampled ROIs, we applied a minimum threshold of 25 vertex pairs for an ROI to be included in a given comparison. Results from dorsal areas falling near this threshold should therefore be interpreted cautiously and treated as preliminary. A natural next step would be to manually delineate these ROIs for each of the 11 subjects, which could yield more accurate, individually tailored boundaries and increase the number of usable vertex pairs in these regions.

Another limitation comes from the relatively small field of view (FOV) of the stimulus display (9°), which constrains the range of stimulus parameters that could be manipulated and analyzed. Within such a restricted FOV, factors such as bar width, sweep speed, and non-overlapping bar movement could not be varied substantially without compromising stimulus coverage, and the resulting mapping stimuli were necessarily more foveally biased than would be optimal for fully characterizing peripheral pRF properties. Future studies using systems with a larger FOV would allow these parameters to be manipulated more freely, providing a more complete picture of how stimulus geometry influences pRF estimates. Additionally, eye-tracking data collected during scanning were not analyzed in the present study; incorporating this information in future work could help verify fixation stability and further constrain interpretations of the foveal bias observed here.

While we employed the established OG and CSS models, future research could further explore these results by implementing other models, such as Difference of Gaussians (DoG) or Divisive Normalization (DN) for pRF fitting. The DN model, in particular, may help clarify the size-dependent effects observed in the present results (differences seen across element sizes 25, 50, and 100) as its additional surround-suppression component could account for some of the effect sizes attributed here to stimulus manipulations, potentially reducing or reshaping these differences when modeled explicitly (Aqil et al., 2021).

The last potential limitation involves the differences in spatial excitation between our stimulus categories. While CB filled the entire moving bar aperture, RW and FF consisted of discrete characters that did not stimulate every point within the bar, particularly at the corners and edges. As our pRF model used a binary mask that only marked the rectangular position of the bar and not the specific character shapes within it, the model assumed a uniform density of excitation that did not exist for RW and FF. This mismatch in stimulus-driven excitation may have influenced the systematic differences in eccentricity we observed between CB and RW. Implementing pixel-wise models that capture exact string distributions could clarify the extent to which spatial border effects underlie the stimulus-dependent variations seen in pRF eccentricity estimates.

## Conclusions

The present study establishes a sensory baseline for the human reading network, bridging low-level visual space and high-level orthographic processing through a dense-sampling fMRI approach. Thre main findings emerge from this work.

First, we provided a robust subject-level replication of hierarchical stimulus-dependent pRF shifts. Our findings confirm that orthographic carriers (words and false-fonts) elicit systematically more foveal eccentricity estimates than checkerboards, demonstrating that this spatial bias is a stable, structural feature of the ventral stream’s architecture.

Second, we characterized the critical role of FWS (Frequency, Width, and Size) parameters in parameter stability. Our analysis revealed that pRF estimates are not neutral readouts but are influenced by temporal and spatial stimulus properties, with lower flickering frequencies and smaller element sizes generally yielding more peripheral estimates. This emphasizes the necessity of matching these low-level features in comparative studies of the reading circuitry.

Third, our model comparison indicates that choice of pRF framework (OG vs. CSS) is a meaningful factor in spatial mapping, as linear models like OG systematically yield more peripheral estimates than non-linear models in higher-order regions.

Finally, we designed and validated the Word-Center (WC) paradigm for decoupling bottom-up sensory bar-sweeps from top-down central cognitive processing. The paradigm demonstrated stimulus equivalence, yielding reliable retinotopic estimates while simultaneously serving as a functional localizer for specialized reading regions such as mOTS, pOTS and OWA.

In conclusion, this work provides a methodological framework and a validated hybrid paradigm. Our future research will utilize the Word-Center paradigm alongside eye-tracking and varying cognitive loads to isolate the specific top-down attentional mechanisms driving these spatial shifts, working towards a multi-signal model of the human reading hierarchy.

## Acknowledgements

We used artificial intelligence-based tools to assist in writing, editing, and reviewing the manuscript. The authors reviewed and approved all content to ensure accuracy and integrity.

